# Pharmacological targeting of EED is an effective therapeutic strategy in cellular models of incurable neuroendocrine prostate cancer

**DOI:** 10.64898/2026.02.24.707688

**Authors:** Kayleigh J. A. Orchard, George Bryant, Maryam Latarani, Isabel R. Misir, Satyanarayana M. Yerra, Christos Velanis, Marta Banchi, Irene Fischetti, Steven L. Turnball, Mark Eccleston, Theresa K. Kelly, Edwina Burke, Zoe R. Maylin, Guido Bocci, Jonathan Shamash, Daniel Berney, Adam Brentnall, Shushuke Akatmatsu, Dong Lin, Yong-Jie Lu, Elena Jachetti, Yuzhuo Wang, Francesco Crea

## Abstract

**Background:** Neuroendocrine Prostate Cancer (NEPC) is an incurable malignancy, originating from the trans-differentiation of prostate adenocarcinoma (PRAD). Compared to PRAD, NEPC shows over-activation of Polycomb Repressive complex-1(PRC1) and-2 (PRC2), which are multiprotein epigenetic writers that drive cancer progression via tumour suppressor gene silencing. Tazemetostat is a PRC2 inhibitor approved for the treatment of sarcomas and lymphomas. ORIC-944 is a novel EED (Embryonic Ectoderm Development) inhibitor, which is being tested in clinical trials. EED is an attractive target as it functions as a key component of both PRC1 and PRC2.

**Objective and Methods:** We compared the anticancer effects of tazemetostat and ORIC-944 in NEPC and PRAD cells. Cells were exposed to various concentrations of the two compounds to measure effects on cell viability (IC_50_) and apoptosis (flow cytometry). PRC2 inhibition was confirmed by measuring histone H3 Lys 27 trimethylation (H3K27me3) via ELISA and Western Blot. RNA Sequencing and pathway analysis was conducted to study modes of actions of tazemetostat vs ORIC-944.

**Results:** Unlike tazemetostat, ORIC-944 causes dose-dependent growth inhibition in both NEPC and PRAD cells. In this context, EED targeting achieves IC_50_ values that are comparable to those of compounds used for the clinical treatment of advanced prostate cancer. Moreover, ORIC-944 (but not tazemetostat) causes significant apoptosis in NEPC cells. Both tazemetostat and ORIC-944 reduce H3K27me3. Mechanistically, both compounds reactivate the expression of known PRC2 targets, such as genes that control neural differentiation. However, the EED inhibitor also reactivates PRC1 targets, including pro-apoptotic and anti-proliferating genes (e.g. metallothionines). This evidence suggests that EED inhibition is a promising therapeutic strategy for NEPC.

## 1. Introduction

When exposed to androgen deprivation therapy (ADT) or androgen receptor pathway inhibitors (ARPIs), approximately 15% of prostatic adenocarcinomas (PRAD) trans-differentiate into neuroendocrine prostate cancers (NEPC)^[1]^. This phenomenon is driven by epigenetically controlled phenotypic plasticity and is facilitated by hypoxia^[2]^. NEPCs are androgen-indifferent and resistant to both hormonal therapies and chemotherapy. NEPC diagnosis entails a dismal prognosis^[3]^. Hence the identification of effective therapeutic strategies for NEPC is an unmet clinical need.

The Polycomb Repressive Complexes 1 and 2 (PRC1, PRC2) are highly conserved epigenetic writers that control lineage specification in mammalian cells^[4,5]^. PRC1- and PRC2-dependent gene silencing are mediated by histone H2a Lys 119 monoubiquitylation (H2aK119ub) and histone H3 Lys 27 trimethylation (H3K27me3), respectively. In several malignancies, PRC1 and PRC2 functions are hijacked to silence tumour suppressor genes and to promote cancer progression^[4,6]^. The histone methyltransferase EZH2 is thought to be the main driver of PRC2-dependent tumour progression^[7]^. For these reasons, a series of selective EZH2 inhibitors has been developed and tested in clinical trials^[8]^. The EZH2 inhibitor tazemetostat is approved for the treatment of epithelioid sarcomas and B-cell lymphomas.

We showed that PRC1 and PRC2 components are highly up-regulated in NEPC, compared to PRAD^[9]^ and that the PRC1 protein CBX2 mediates NEPC progression^[10]^. Both NMYC^[11]^ and WNT pathway drive NEPC trans-differentiation via EZH2^[12]^. Hence it is conceivable that PRC1 and PRC2 inhibitors could work as anticancer agents in NEPC. However, pharmacological inhibition of EZH2 does not significantly affect the growth of NEPC cells^[13, 14]^, whilst retaining *in vitro* and *in vivo* anticancer activity in PRAD ^[15]^. Potent small molecule inhibitors of EED have been recently developed^[16]^. EED has a double function: it is required for PRC2 anchorage to H3K27 residues and for PRC1-mediated H2aK119ub^[17]^. Hence EED targeting compounds could inhibit both PRC1 and PRC2.

Here we compared the pharmacological and epigenetic effects of a traditional EZH2 inhibitor (tazemetostat) vs the new EED inhibitor ORIC-944. Our results show that ORIC-944 is effective both in PRAD and NEPC cells, and that EED inhibition elicits a gene expression profile that is partially different from EZH2 inhibition and potentially associated to the reactivation of PRC1-specific genes. This observation could explain the different efficacy of EED and EZH2 inhibitors in NEPC cells.

## 2. Materials and Methods

### 2.1 Cell Culture

The LNCaP cell line was purchased from ATCC (CRL-1740**™**), and KUCaP13 were provided by Kyoto University Graduate School of Medicine^[14]^. Both cell lines were cultured in RPMI 1640 medium (Gibco; 21875034) supplemented with 10% heat-inactivated fetal bovine serum (Gibco; A5670701), 6.25ml 100mM HEPES (KUCaP13 (Gibco; 15630056)) and 1% penicillin-streptomycin (Gibco; 15140122). ST4787, a murine model of NEPC, and T23, a murine model of adenocarcinoma, were provided by Istituto Nazionale dei Tumori^[18,19]^ and were cultured in DMEM medium (Gibco; 11995065) supplemented with 10% heat-inactivated fetal bovine serum, 5ml 100mM HEPES, 5ml 10mM Sodium Pyruvate (Gibco; 11360070) and 1% penicillin-streptomycin. For experiments investigating the effect of hypoxia on cell viability and pharmacological response, cultures were moved into the hypoxia chamber (Baker Ruskinn) after trypsinization, and grown at 37°C, 2% O_2_ for two days before experiments were initiated. Media, including Trypsin-EDTA (0.25%, Gibco; 25200072) and plastics, were also moved to the hypoxia chamber or PhoxBox (Baker Ruskinn) for two days prior to experimentation.

### 2.2 Cell Viability and Dose Response

Cell lines were treated with varying concentrations (25-, 10-, 1-, 0.1-, 0.01, and 0.001µM) of EZH2i tazemetostat (MedChemExpress; HY-13803), or EEDi ORIC-944 (ORIC Pharmaceuticals Inc, MedChemExpress; HY-158102), and 0.05% DMSO (vehicle control (MedChemExpress; HY-Y0320)) for seven days in both normoxia (21% O_2_) and hypoxia (2% O_2_). The seven-day experiment involved single-dose drug treatment for four days, media change and drug replenishment on day four and then incubation until day seven, and this is consistent throughout experiments where seven days of treatment has been used (See Section 2.3 and 2.5). In normoxia, cell viability was measured using Cell Titre Blue (Promega; G8080) which is a fluorescent method based on the conversion of resazurin to resorufin by viable cells. In hypoxia, cell viability was determined using an automated cell counter (ThermoFisher; Countess™ 3), and trypan blue (0.4%, Gibco; 15250061) in a 1:1 ratio. Using nonlinear regression of log(inhibitor) vs. response, dose-response curves were produced which allowed for the identification of the half-maximal inhibitory concentration (IC_50_) (GraphPad Prism 10 (10.6.1)).

### 2.3 H3K27me3 quantification

#### 2.3.1 Histone Extraction, Quantification and ELISA

Histones from KUCaP13, LNCaP, ST4787 and T23 were extracted after a seven-day treatment with 5µM, 1µM EZH2i or EEDi and DMSO (0.05%) using Histone Extraction Kit (Abcam; ab113476) following the manufacturers protocol. Extracted histones were quantified using the Pierce™ Dilution Free™ Rapid Gold BCA Protein Assay (ThermoFisher; A55860) and were subsequently used in the Active Motif ELISA assay for Total Histone H3 and H3K27me3 following manufacturers protocol (Active Motif; 53110 and 53106). Quantification of Total H3 and H3K27me3 in cell lines after the aforementioned treatment with EZH2i or EEDi was achieved by producing a standard curve using either recombinant histone H3 protein, or recombinant H3K27me3 protein as suggested by the manufacturer using GraphPad Prism 10 (10.6.1). Finally, following the manufacturer’s protocol, the methylation of samples at H3K27me3 was normalised to Total H3.

#### 2.3.2 Western Blot

Extracted histones were quantified using the Pierce™ Dilution Free™ Rapid Gold BCA Protein Assay (ThermoFisher; A55860) and based on the predicted molecular weight of the protein of interest, gel electrophoresis was performed on Novex Tris-Glycine 10-20% WedgeWell protein gels (ThermoFisher Scientific; XP10205BOX). Proteins were transferred to Amersham™ 0.2µm nitrocellulose membranes (Cytiva; 10600019) and blocked with 5% milk or 5% bovine serum albumin (Merck; A9418) prepared in tris-buffered saline with 0.1% Tween 20 (ThermoFisher Scientific; J77500.K2). Primary antibodies against H3K27me3 (Cell Signalling Technology; 9733S; 1:1000 dilution), H2AK119ub (Cell Signalling Technology; 8240Sl 1:1000 dilution) and Total H3 (Cell Signalling Technology; 9715S; 1:1000 dilution) were incubated overnight at 4°C. After washing, the membrane was then incubated with the HRP-conjugated secondary antibody (ThermoFisher Scientific; 31460) for 1 hour at room temperature (RT). Detection was performed using Amersham™ ECL™ Western Blotting Reagents (Cytiva; RPN2106) and membranes were visualised using the G:BOX Chemi XX6 (Syngene).

### 2.4 Apoptosis

#### 2.4.1 Cell Preparation

ST4787 cells were grown in 6-well plates and maintained at 37°C, 21% O_2_ and 5% CO_2_ for 24 hours, when cells were treated with either DMSO (0.1%), ORIC-944, tazemetostat and carboplatin in concentrations of 10-, 5- and 1µM for 3 days. After treatment cells were detached from the plates and collected in sample tubes along with the supernatant. The cell suspension was then stained using eBioscience™ Annexin V Apoptosis Detection Kit: Annexin V conjugated to Allophycocyanin (AnV-APC) and propidium iodide (PI) (ThermoFisher Scientific; 88-8007-74) following the manufacturers protocol. PI is a viability dye that binds to DNA and will only enter cells with compromised membranes. AnV-APC is a calcium dependent protein that binds to phosphatidylserine (PS). In healthy cells, PS is a membrane bound protein on the inner envelope of the membrane. During apoptosis PS moves to the outer envelope of the cell membrane, allowing for the binding of AnV-APC.

#### 2.4.2 Flow Cytometry

Stained cell suspensions were transferred to BD Falcon™ 5mL polystyrene round-bottom tubes (Corning; 352052) and analysed using the Novocyte 3000 3 Laser Flow Cytometer (Aligent) with an autosampler. Depending on which limit was reached first, either 50,000 events or 100µl of sample were analysed. Data was plotted on an FSC-H x SSC-H plot, gated to isolate the cell population from noise and then using FSC-H x FSC-A to gate single cells. Finally, the data were plotted using APC-H x PE Texas red-H to identify apoptotic and necrotic cell populations and gated using a quadrant gate. Stained controls were used to set gating and consisted of heat shocked cells to induce apoptosis and necrosis and untreated cells. These were then divided and stained as no stain, AnV-APC only, PI only and AnV-APC and PI. Once the gating was set on the control groups it was applied to the experimental groups, and the resulting population distribution was used for statistical analysis using GraphPad Prism 10 (10.6.1).

### 2.5 RNA-Sequencing and Analysis

Both KUCaP13 and ST4787 were expanded in culture and treated with either 5µM EZH2i tazemetostat, 5µM EEDi ORIC-944, DMSO (0.05% vehicle control), or untreated, for seven days to obtain a minimum of 2.5 x 10^5^ cells for bulk RNA-Seq. Cells were then pelleted and snap-frozen following Active Motifs protocol for sample preparation. Three biological replicates from both experimental and control groups were submitted for analysis, and all downstream sample preparation and analysis was conducted by Active Motif. RNA was analysed using an RNA ScreenTape Assay for TapeStation (Aligent) and samples with an RNA integrity number (RIN) of 9.7 – 10 were used for library preparation using the Illumina TruSeq RNA Sample Preparation v2 guide (Illumina). Libraries were sequenced on an Illumina NextSeq 500 as paired-end 150 bp reads (PE150) and reads were mapped to either the human genome (hg38) or mouse genome (mm39) using the STAR (v2.6.1)^[20]^ algorithm with default settings. Salmon^[21]^ was then used to count the number of fragments overlapping predefined genes, and after obtaining gene expression count matrixes, differential analysis was performed to identify statistically significant differential genes using DESeq2 (v1.28.0)^[22]^ with a p. adjusted value of less than 0.05, and a log fold change of 1. Finally, gene set enrichment analysis (GSEA) to identify biological pathways, gene ontology terms or other functional categories that are enriched in the genes of interest was performed using clusterProfiler (v4.8.2)^[23]^ with default parameters.

## 3. Results

### EED targeting induces dose-dependent growth inhibition in PRAD and NEPC

To compare the effects of EZH2 and EED inhibition, we exposed PRAD and NEPC cells to different concentrations of tazemetostat and ORIC-944. As shown in Figure 1, A - C tazemetostat induced dose-dependent growth inhibition in LNCaP cells (PRAD) with an IC_50_ of 0.052±0.011µM, but was ineffective in all the other cell lines, where we could not calculate an IC_50_. However, ORIC-944 caused dose-dependent growth inhibition in both NEPC and PRAD cells, with IC_50_ values ranging from 0.13 to 2.84 µM (Table S1). ORIC-944 remained effective in NEPC cells grown under hypoxia, which is a known mediator of treatment-resistance^[24]^ (Figure 1, D, E). Notably, in ST4787 NEPC cells, ORIC-944 was at least as effective as some compounds in clinical use for advanced prostate cancer treatment, such as carboplatin and olaparib (Figure 1 F, G and Table S1).

**Figure 1.**
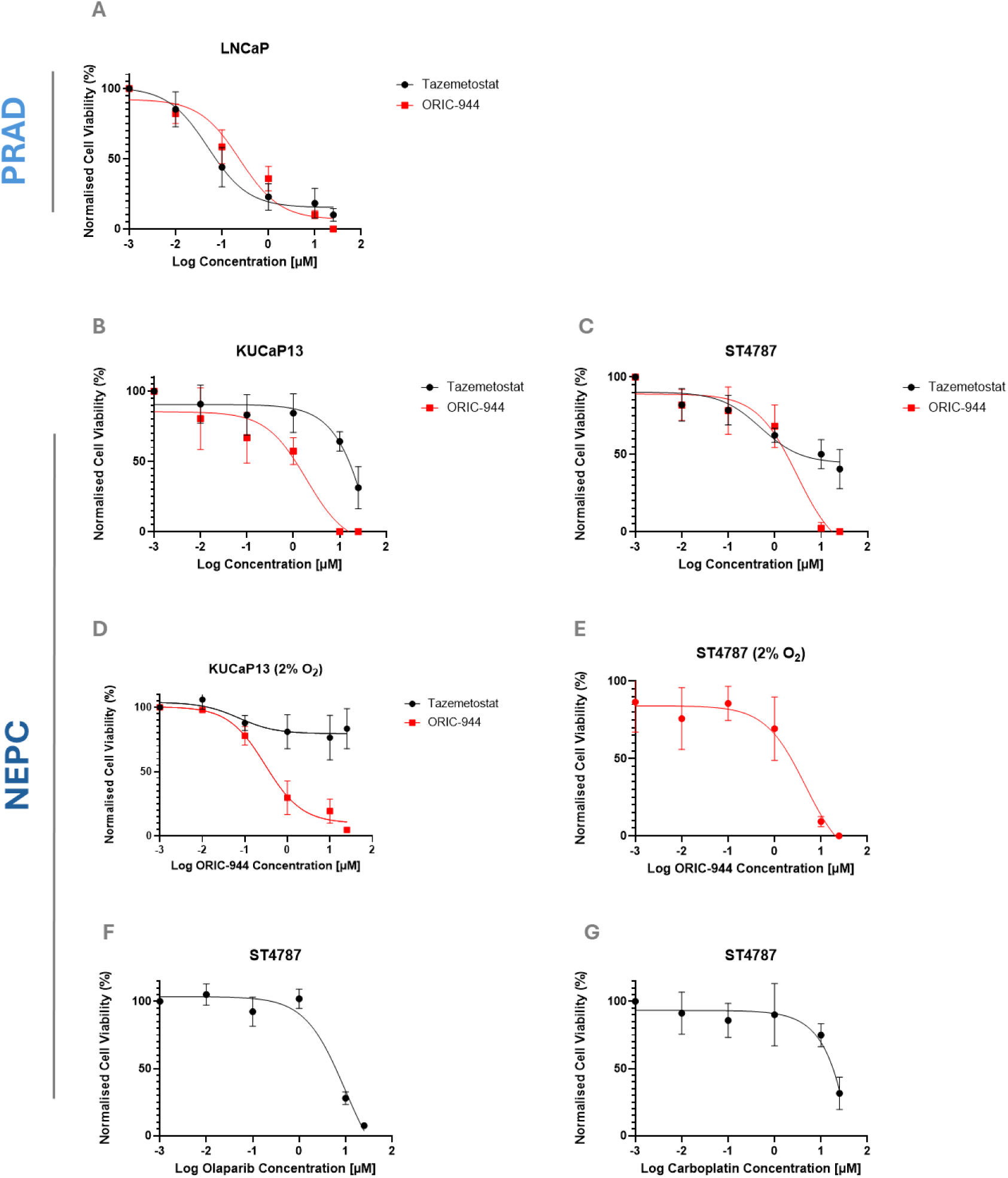
Growth inhibitory effects of PRC2 inhibitors and clinically used compounds in PRAD and NEPC cells. Cells were treated with 25µM – 0.001µM of ORIC-944 or tazemetostat in normoxia (21% O_2_), and Cell Titre Blue® viability assay was performed after 7 days of treatment. The same treatment and analysis was repeated in hypoxia (2% O_2_) for ORIC-944 and trypan blue was used to determine viability after 7 days (**D – E**). NEPC model ST4787 were also treated with 25µM – 0.001µM of either olaparib (**F**) and carboplatin (**G**) for 7 days, and Cell Titre Blue® viability assay was performed. All plots show average of 3 independent experiments with lines representing standard deviations.

To investigate whether the efficacy of ORIC-944 could be attributed to double targeting of EZH1- and EZH2- containing PRC2, we exposed NEPC cells to the double EZH1/EZH2 inhibitor valemetostat (Figure S1). We could not calculate the IC_50_ for this compound in ST4787 cells, whilst in KucaP13 the IC_50_ was higher than 10 µM. This shows that ORIC-944 is effective in both NEPC and PRAD cells.

### EED inhibition causes apoptosis in NEPC cells

To investigate the molecular correlates of drug activity, we measured total H3K27me3 levels in cells exposed to ORIC-944, tazemetostat or control (DMSO). As expected H3K27me3 levels were consistently reduced by both drugs (Figure S2). However, based on flow cytometry analysis only ORIC-944 caused significant cell number reduction and induced apoptosis in NEPC cells (Figure 2). Notably, this apoptotic effect was evident after 3 days of treatment. Taken together, these results suggest that ORIC-944, but not tazemetostat, inhibits the proliferation of NEPC cells, and that this effect is mediated by apoptosis.

**Figure 2.**
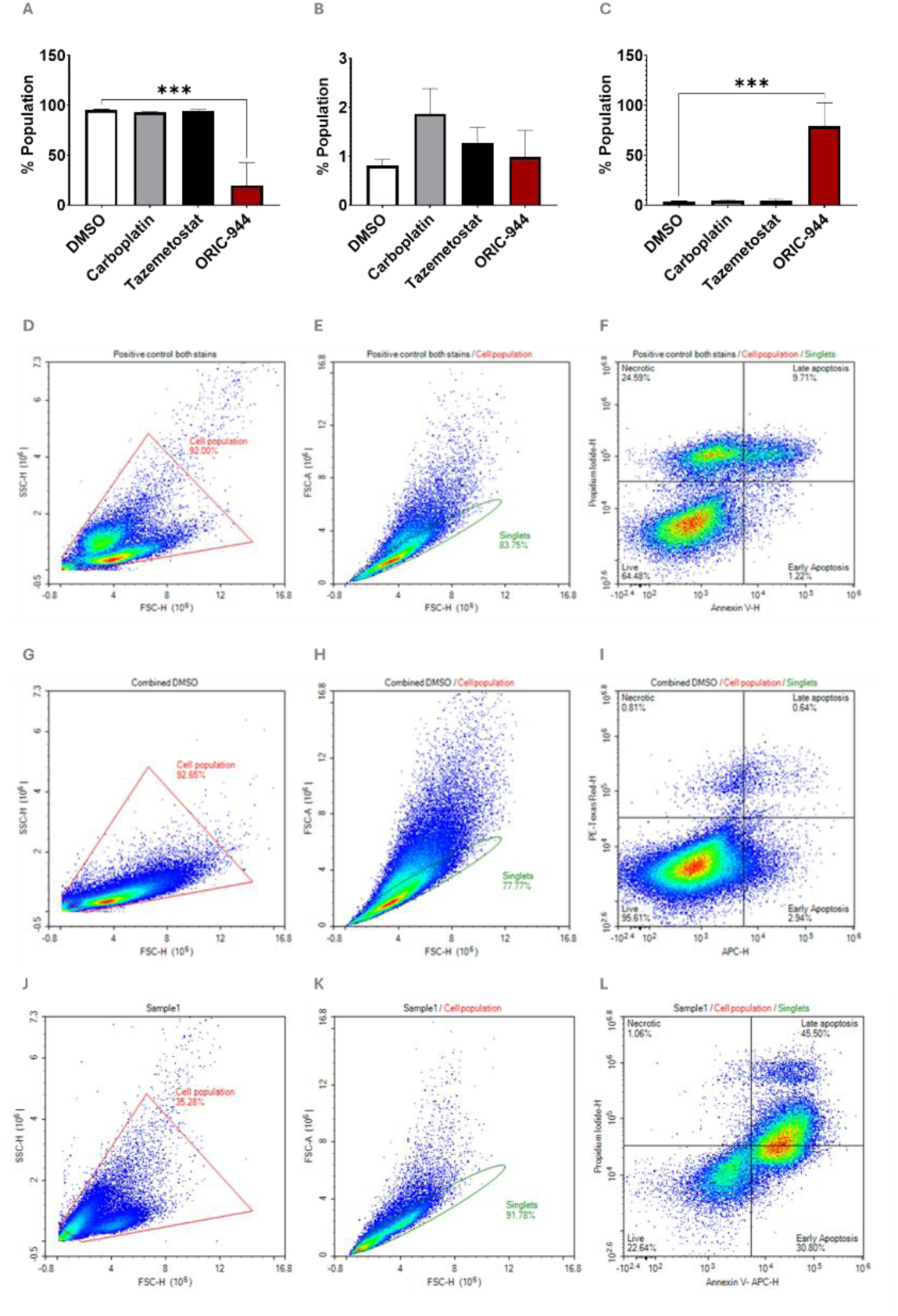
Flow cytometry analysis of viability and apoptosis in NEPC cells exposed to PRC2 inhibitors or carboplatin. (**A – C**) Bar plots showing results from flow cytometry analysis in ST4787 cells after 3 days of treatment with DMSO (vehicle), carboplatin, ORIC-944 (10µM) and tazemetostat (10µM). A: cell viability; B: necrosis; C: apoptosis. Bars show average of 3 independent experiments; lines represent standard deviation and the statistical analysis used was a one-way ANOVA (Tukey’s multiple comparisons) and statistical pairwise comparisons are overlayed (p***<0.0005). **D-L:** Representative scatter plots from flow cytometer analysis of cells treated with positive control (heat shock, D - F), DMSO (G - I) and ORIC-944 (J - L). Forwards Scatter Height (FSC-H- D, G, J) x Side Scatter Height (SSC-H) identifies cell population. Colours indicate numerosity of cell populations, red = more numerous, blue = less numerous. FSC-H x Forwards Scatter Area (FSC-A, **E, H, K**) based on cell population gating, identifies the single cell population. Annexin V intensity (H) x Propidium Iodide intensity (H) plot based on gating from singlets (**F, I, L**), with quadrant gating identifying live, necrotic, late apoptotic and early apoptotic cells.

### Both EED and EZH2 inhibitors activate neural-specific genes in NEPC cells

To further dissect the genetic networks elicited by EED and EZH2 inhibition in NEPC cells, and to identify potential mechanisms of resistance, we analysed the transcriptome of NEPC cells exposed to tazemetostat, ORIC-944 or DMSO for 7 days. This analysis revealed that both PRC inhibitors predominantly increased gene expression, in line with the suppression of canonical Polycomb functions (Figure 3, A, B (KUCaP13) and E, F (ST4787). The gene ontology of up-regulated transcripts showed that both PRC inhibitors induced the activation of neural differentiation and cell specification networks, with substantial overlap of pathways between compounds and between the two cell lines (Figure 3, C, D (KUCaP13) and G, H (ST4787). Upon treatment with tazemetostat and ORIC-944, known Polycomb target genes that modulate the WNT pathway^[25]^ were significantly up-regulated in both KUCaP13 and ST4787 cells (Figure 3, I - J). As previously described^[13]^, both PRC inhibitors increased the transcription of genes that contain bivalent promoters, such as Human Leucocyte Antigen coding *loci* (Figure 3, H).

**Figure 3.**
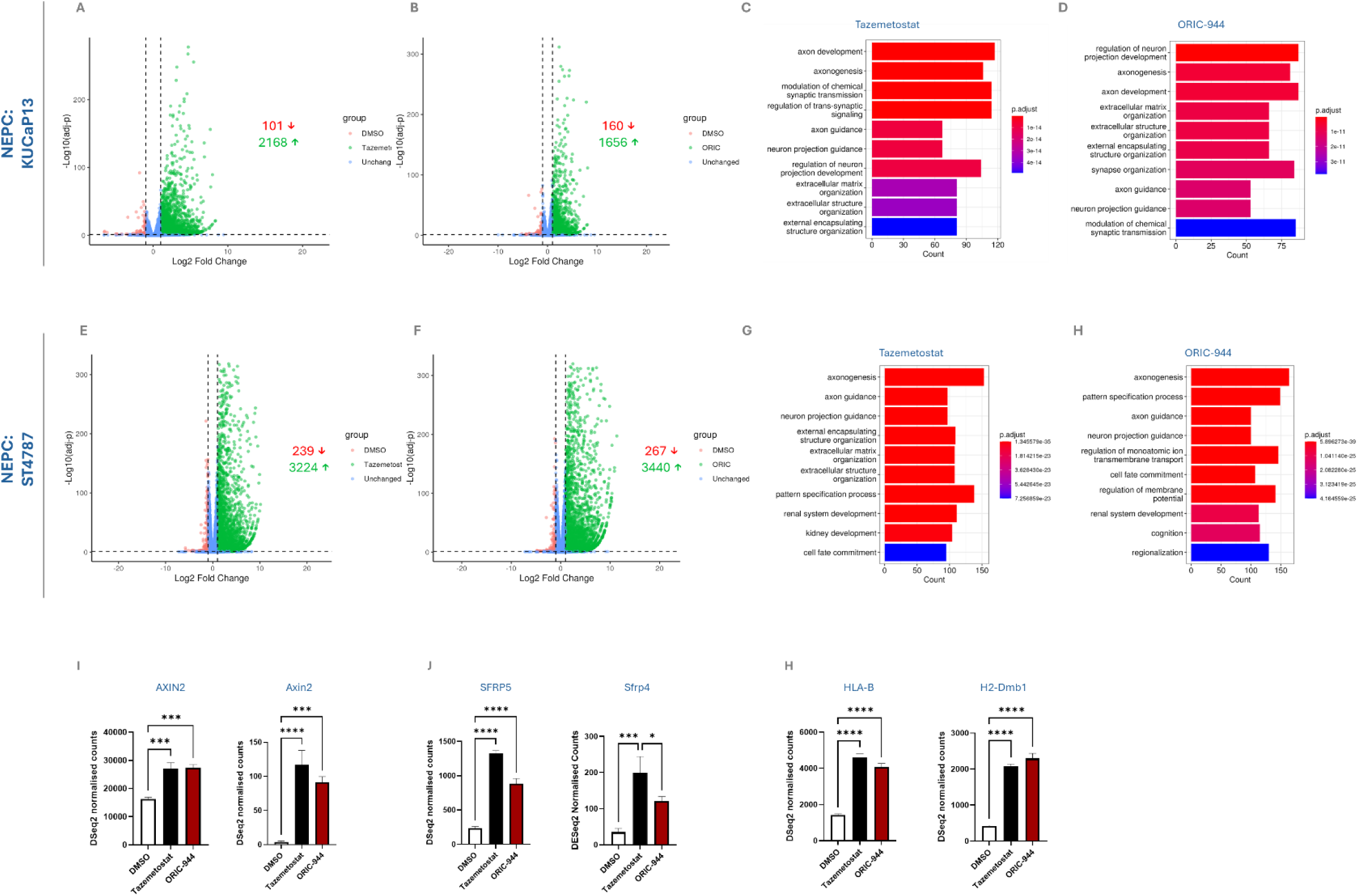
RNA Sequencing reveals shared pathways activated by different PRC2 inhibitor. Volcano plots showing upregulated (green), unchanged (blue) and downregulated (red) genes upon treatment with tazemetostat or ORIC-944 in NEPC models – KUCaP13 (**A, B**) and ST4787 (**E, F**). Arrows and colours represent the number of genes significantly upregulated (↑ - green) and downregulated (↓ - red) in each dataset. Cut-off values: log2FoldChange >1 (upregulation), or <1 (downregulation), and a p.adj.<0.05. Gene ontology analysis of significant differentially expressed genes (tazemetostat vs DMSO (**C**), ORIC vs DMSO (**D**) in KUCaP13, and ST4787 (**G, H**). Expression of two known Polycomb targets (**I, J**) upon treatment with tazemetostat vs DMSO in KUCaP13 (left) and ST4787 (right). Expression of two HLA genes upon treatment with tazemetostat vs DMSO in KUCaP13 (left) and ST4787 (right) (**H – K**). Each p value is calculated by ANOVA Dunnett’s post-hoc test ***p<0.001; ****p<0.0001. Bars represent average values of 3 independent experiments; lines represent standard deviation.

### A subset of proapoptotic genes is selectively activated by EED inhibition

Given the phenotypic differences elicited by EZH2 vs EED inhibition, we investigated whether we could identify a subset of transcripts that were preferentially activated by ORIC-944 vs tazemetostat (Preferentially ORIC-944 Dependent genes-PODs). By applying a Log2 Fold change>1 and a padj<0.05 (ORIC-944 vs tazemetostat), we found 10 and 362 PODs in KUCaP13 and ST4787 cells, respectively (Table S2 and S3). Gene ontology analysis revealed that in KUCaP13 cells, which are the most sensitive to ORIC-944, PODs were enriched for “negative regulation of growth” (Figure 4, A). This ontology included three metallothionines, some of which are known Polycomb targets^[26]^ (Figure 4, B – D). In the ST4787 cells, PODs were enriched for several developmental pathways, especially “pattern specification” and “regionalization” (Figure 4, E). This ontology included both known tumour suppressor genes and mediators of the Wnt pathway (Figure 4, F - H). We then crossed our list of PODs with genes that were identified as selective PRC1 targets, selective PRC2 targets, or targets of both complexes in prostate cancer cells^[17]^. Notably, PODs were enriched for selective PRC1 targets, both in KUCaP13 and in ST4787 cells (Figure 4, I, J and Table S2 and S3).

**Figure 4.**
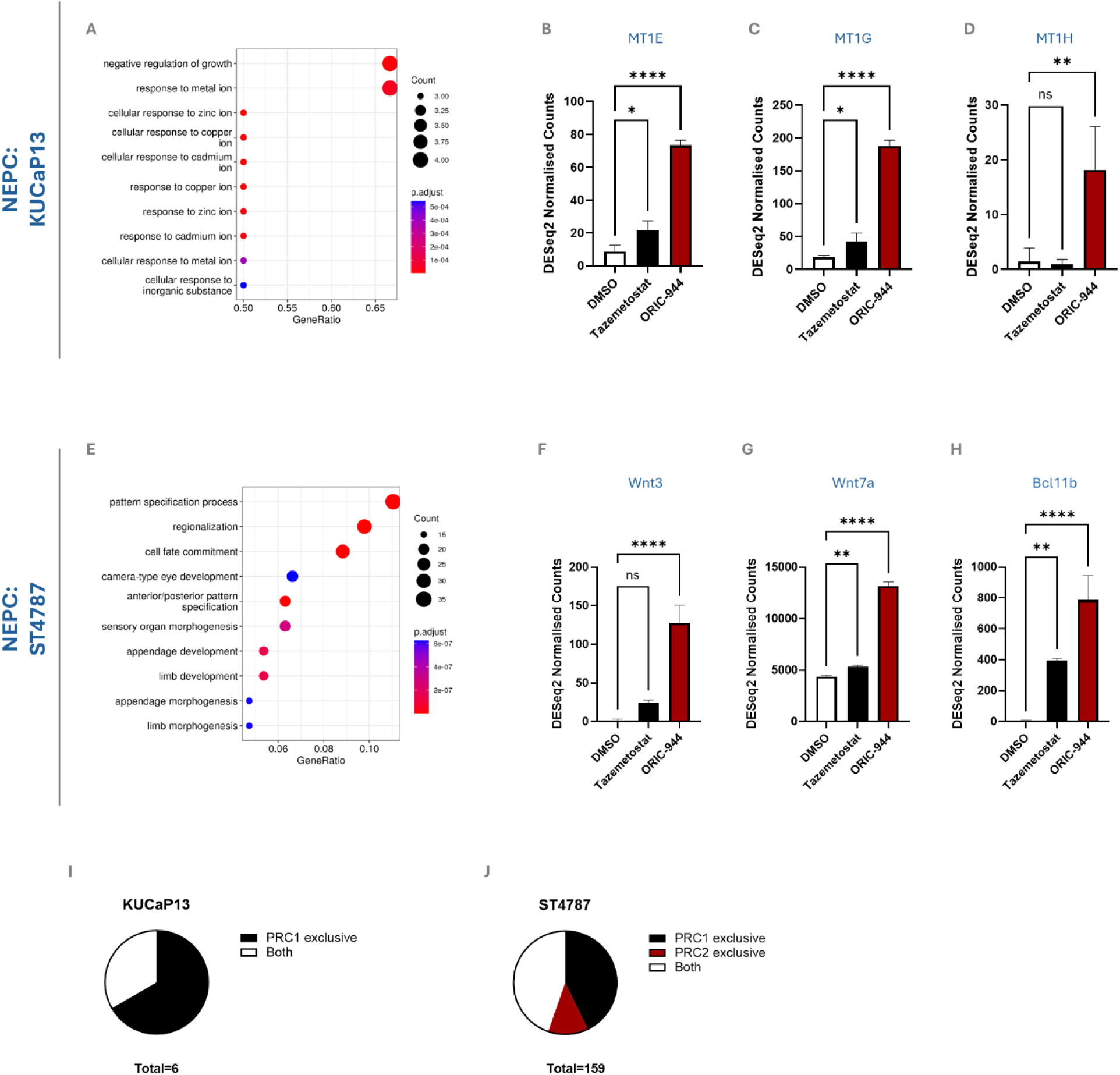
Identification of Preferentially ORIC-944 Dependent genes (PODs). Gene Ontology analysis of genes differentially expressed upon treatment with ORIC-944 in KUCaP13 (**A**). Expression of three genes associated with the most upregulated pathway (negative regulation of growth) in KUCaP13 (**B – D**). Gene Ontology analysis of genes differentially expressed upon treatment with ORIC-944 in ST4787 (**E**). Expression of three PODs in ST4787 (**F – H**). B – D and F – H p values were calculated by ANOVA Dunnett’s post-hoc test ***p<0.001; ****p<0.0001. Bars represent average values of 3 independent experiments; lines represent standard deviation. Percentage of PODs that are PRC1-selective targets, PRC2-selective targets or shared targets in KUCaP13 and ST4787 cells (**I – J**; See also Table S2 and S3).

## Discussion

In this study, we showed that the EED-targeting compound ORIC-944 causes dose-dependent growth inhibition and apoptosis in NEPC cells. Notably, ORIC-944 is much more effective than both tazemetostat and carboplatin at inducing apoptosis in NEPC cells (Figure 2).

Despite the prominent pathogenetic role of PRC2 in most stages of prostate cancer initiation and progression, EZH2 inhibitors were shown to be effective in PRAD, but not in NEPC cells^[15]^. This result seems contrary to the significant up-regulation of both EZH2 and EED in NEPC vs PRAD clinical samples^[13]^. We therefore decided to investigate whether EED inhibition elicited anticancer effects in NEPC.

By inhibiting PRC2 canonical functions, tazemetostat reactivates lineage specific *loci* and genes with bivalent promoters in both PRAD and NEPC^[15]^. However, this phenomenon does not induce growth inhibition and/or apoptosis. Instead, the efficacy of tazemetostat in PRAD is at least in part mediated by AR-EZH2 interactions. Aside from its canonical role, EZH2 can act as an AR co-activator in PRAD cells, thereby increasing the expression of cyclins and other proteins that promote cell proliferation. EZH2 inhibitors can target non-canonical functions of this protein and via this mechanism these compounds reduce cyclin expression. Since NEPC are AR-indifferent and often AR-negative, the inhibition of this PRC2-independent function has no effect on this cancer subtype.

Compared to EZH2 inhibitors, which selectively target PRC2, EED inhibitors offer potential advantages. EED coordinates the sequential activation of PRC2 and PRC1, thereby controlling both H3K27me3 and H2K119aub^[6,17]^. Hence EED inhibitors could simultaneously target both PRC1- and PRC2- dependent pathways. Since both PRCs have been implicated in NEPC pathogenesis EED inhibitors may offer a broader spectrum of epigenetic activity for NEPC therapy.

The two NEPC models employed in this study are AR-indifferent^[14,15]^. In line with previous evidence, we found that tazemetostat is not effective in KUCaP13 and ST4787 cells, whilst causing dose-dependent growth inhibition in PRAD cells. As expected, both EED and EZH2 inhibitors cause the activation of neural lineage specific pathways, and of genes putatively associated with bivalent promoters. However, the activation of these genetic networks by both compounds cannot explain their different anticancer activity. For this reason, we actively searched for genes that were selectively up-regulated upon ORIC-944, and not upon tazemetostat treatment (PODs). This analysis revealed for the first time that, along with shared genetic networks, EED inhibition activates a subset of genes in both NEPC cell lines. Interestingly, PODs are enriched for growth inhibiting and pro-apoptotic genes. For example, MT1G, MT1H and MT1F are known for promoting apoptosis in several malignancies, including prostate cancer^[27]^. In prostate cancer, MT1G downregulation mediated by promoter hypermethylation contributes to carcinogenesis and poor prognosis. In thyroid cancer, MT1G acts as a tumour suppressor, with restored expression inhibiting cell growth and invasion whilst inducing cell cycle arrest and apoptosis^[28]^. In prostate cancer, MT1G overexpression inhibits cell growth *in vivo* and *vitro*, modulates immune cell infiltration, and reduces immunosuppressive activity to regulate the tumour microenvironment^[29]^. MT1H is highly methylated and downregulated in prostate cancer, and its restored expression suppresses invasion, colony formation, and cell growth by reducing entry into the S and M phases of the cell cycle^[30]^. Downregulation of MT1F in prostate cancer is linked to perineural invasion^[31]^, though its role in prostate cancer, including NEPC, remains unclear. In colorectal cancer, MT1F increases apoptosis, inhibits growth, and reduces migration, invasion and adhesion^[32]^. Therefore, it is conceivable that EED inhibitors re-activate a unique subset of genes that cause apoptosis in NEPC cells. In ST4787, PODs were also enriched for Wnt pathway modulators and pro-apoptotic factors such as Bcl11b^[33]^. In most cases, the Wnt pathway promotes cancer cell survival^[34]^ so our observation may point to a potential resistance mechanism, which we are currently investigating. Mechanistically, PODs are enriched for PRC1 selective targets in both cell lines, confirming the potentially broader epigenetic spectrum of EED inhibitors, compared to EZH2 inhibitors.

In future work, we plan to investigate these potential mechanisms of action. We also plan to confirm the activity of EED inhibitors *in vivo* and to leverage our transcriptomic analyses to identify active epigenetic combination therapies for NEPC.

## Conflicts of Interest

M.E is CEO of ValiRx plc. The other authors are not aware of any affiliations, memberships, funding or financial holdings that may affect the objectivity of this research manuscript.

## Funding

This study was funded by a Prostate Cancer UK grant RIA22-ST2-006, which was granted to Prof. Francesco Crea, The Open University, Milton Keynes in 2023.

## Supplementary Figures

**Figure S1.**
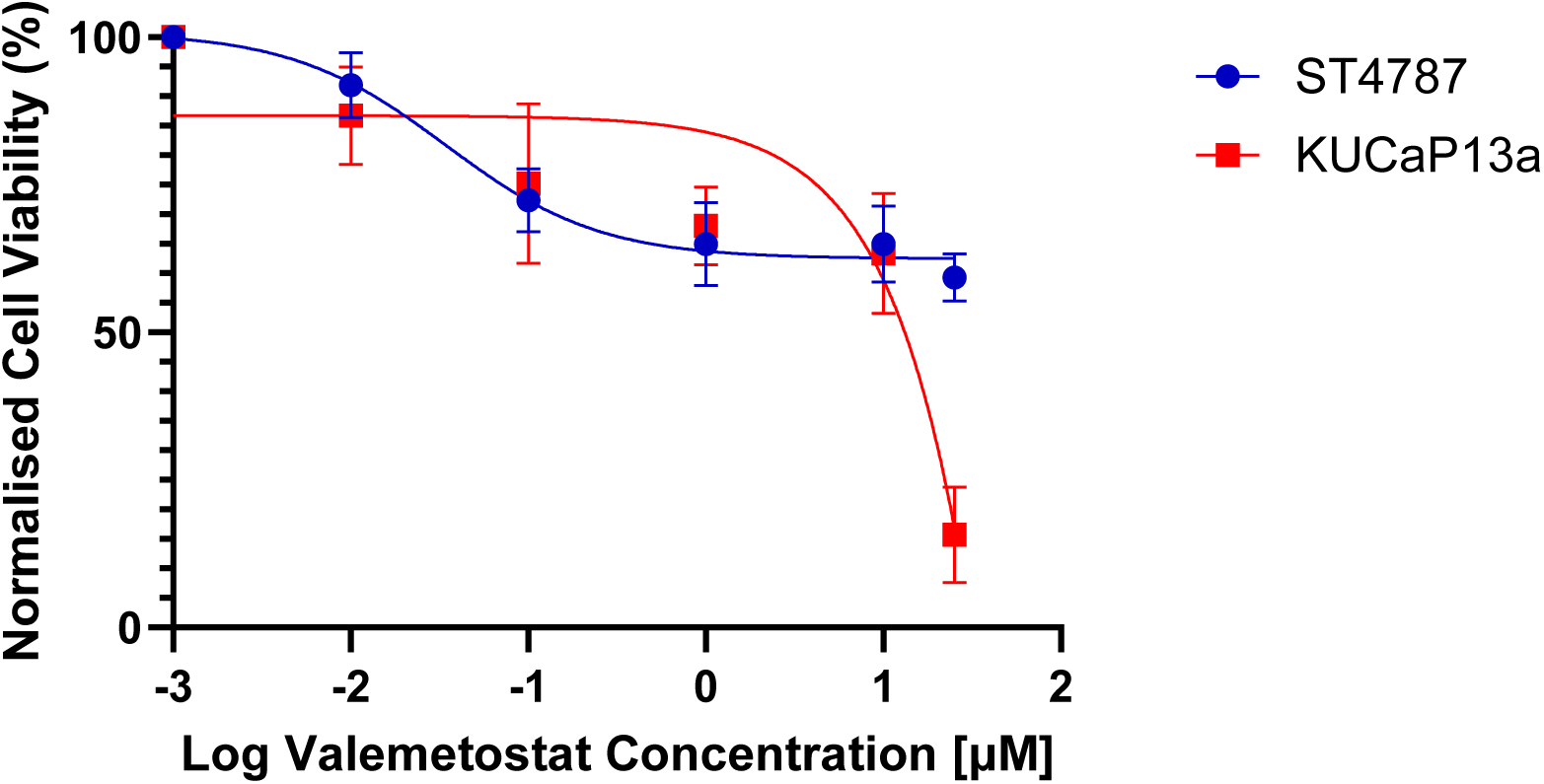
Viability of two NEPC cells exposed to different concentrations of valemetostat. Experiments were performed as described in Figure 1.

**Figure S2.**
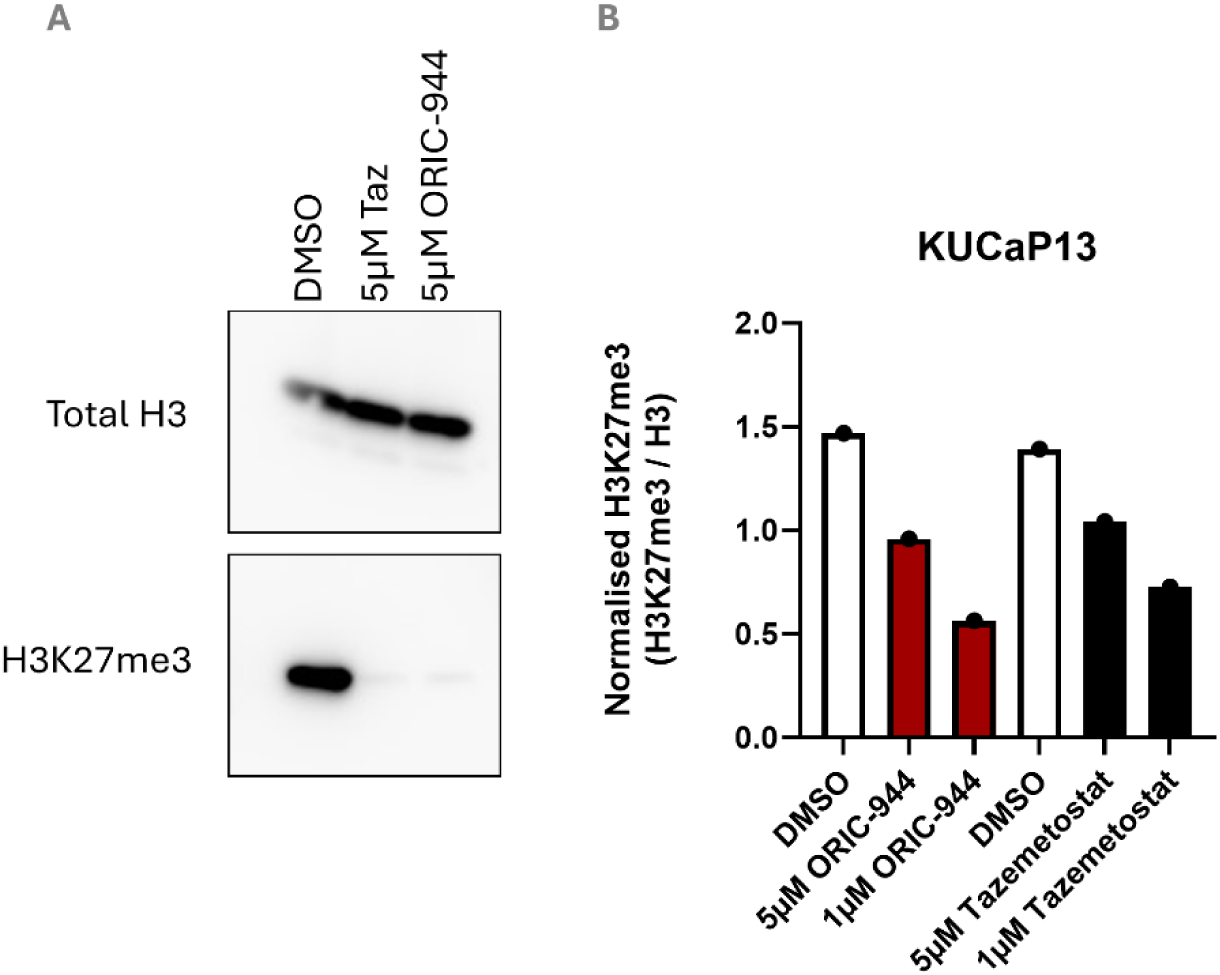
Western blot analysis in KUCaP13 (**A**) confirming the downregulation of H3K27me3 upon tazemetostat (5µM Taz) and ORIC-944 treatment in vitro. Downregulation of H3K27me3 was confirmed via normalisation to Total H3 (H3K27me3 / H3) using ELISA (**B**).

## Supplementary Tables

**Table S1:**
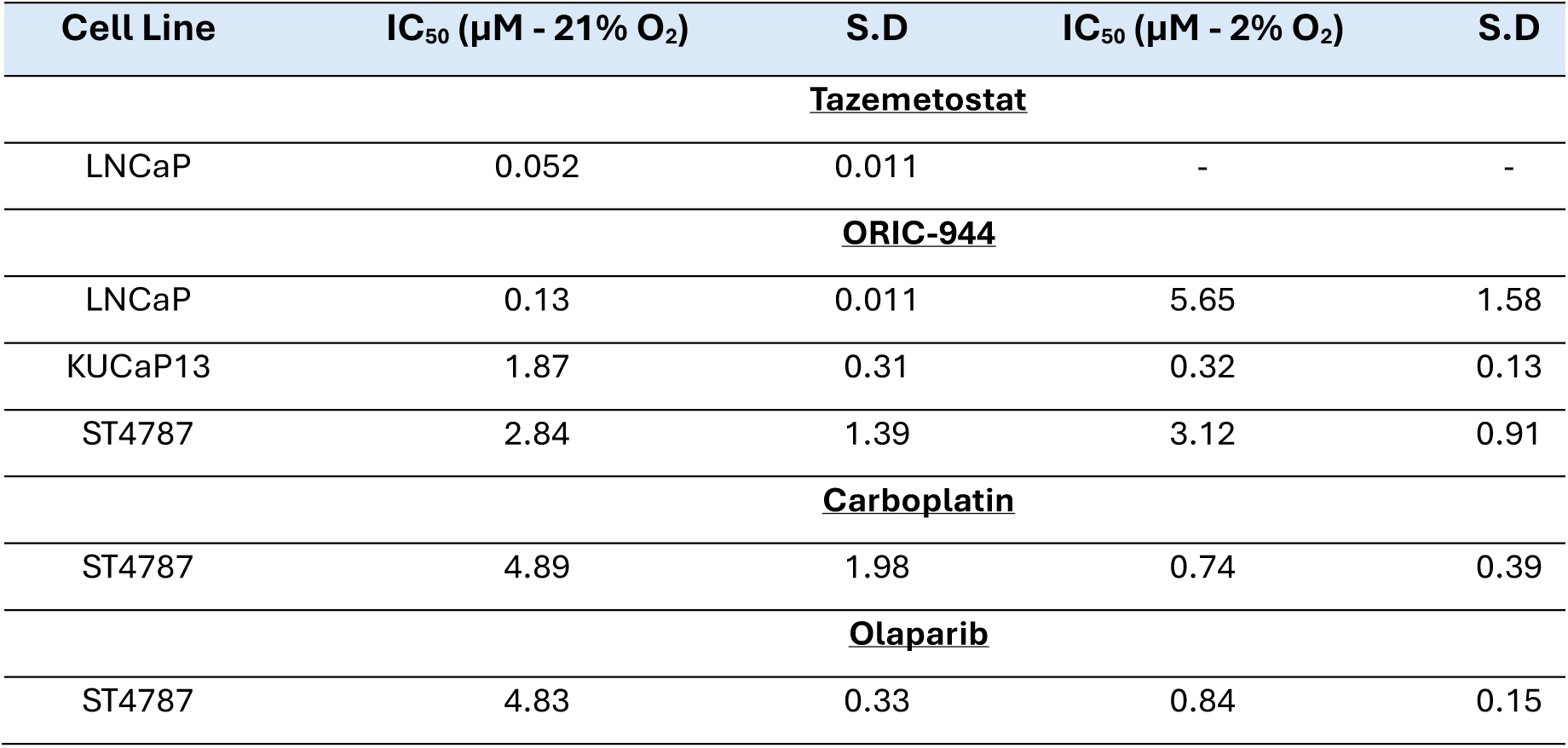
IC_50_ obtained from 7-day incubation with either tazemetostat or ORIC-944 under normoxia (21% O_2_) and hypoxia (2% O_2_) or 7-day incubation with carboplatin or olaparib in normoxia (21% O_2_) and hypoxia (2% O_2_) IC_50_ values (µM) obtained after 7-day incubation with either tazemetostat or ORIC-944, as well as carboplatin or olaparib, and corresponding standard deviation (S.D) values.

| Cell Line | IC<sub>50</sub> (μM - 21% O<sub>2</sub>) | S.D | IC<sub>50</sub> (μM - 2% O<sub>2</sub>) | S.D |
| --- | --- | --- | --- | --- |
| **<u>Tazemetostat</u>** |  |  |  |  |
| LNCaP | 0.052 | 0.011 | - | - |
| **<u>ORIC-944</u>** |  |  |  |  |
| LNCaP | 0.13 | 0.011 | 5.65 | 1.58 |
| KUCaP13 | 1.87 | 0.31 | 0.32 | 0.13 |
| ST4787 | 2.84 | 1.39 | 3.12 | 0.91 |
| **<u>Carboplatin</u>** |  |  |  |  |
| ST4787 | 4.89 | 1.98 | 0.74 | 0.39 |
| **<u>Olaparib</u>** |  |  |  |  |
| ST4787 | 4.83 | 0.33 | 0.84 | 0.15 |

**Table S2:**
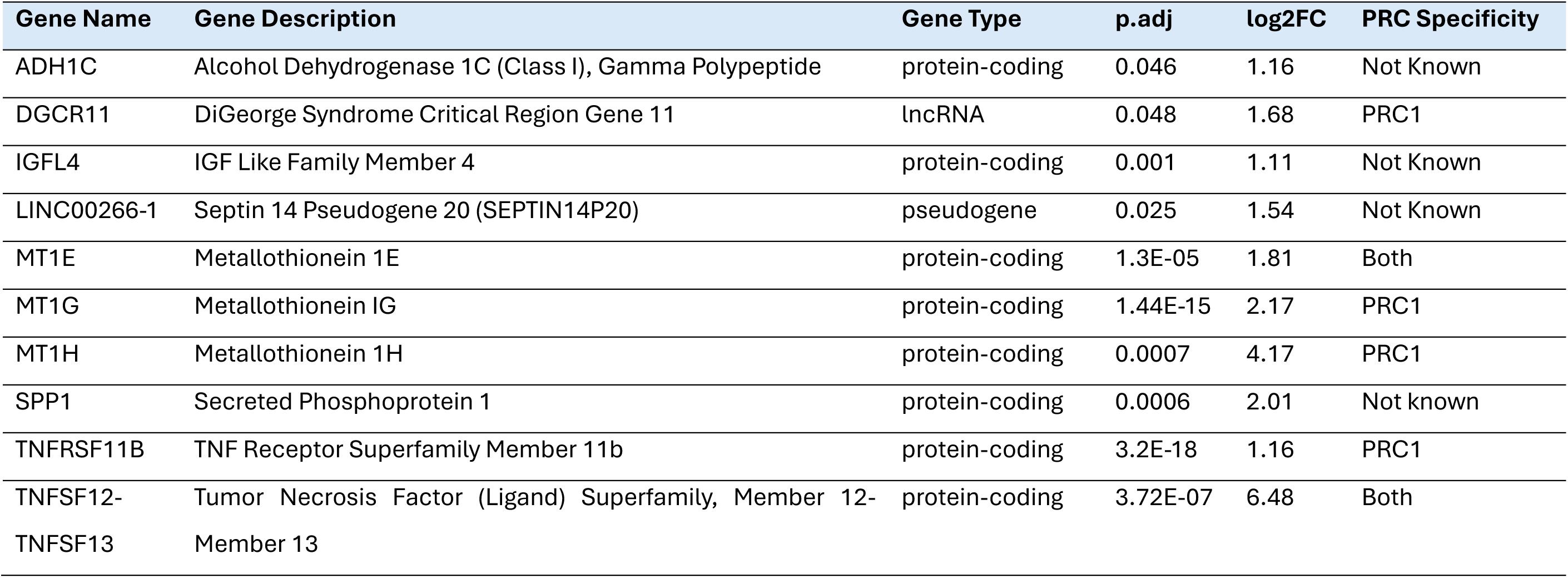
Preferentially ORIC-944 Dependent Genes (PODs) identified from DESeq2 analysis of KUCaP13. DESeq2 analysis of differential gene expression between tazemetostat and ORIC-944 in KUCap13 identified 10 genes that were preferentially ORIC-944 dependent genes (PODs) as well as their p adjusted value (p.adj) and log2FoldChange (log2FC). This identified 8 protein coding, 1 lncRNA and 1 pseudogene. Additionally, using data obtained from DU145 scramble cells^[33]^, out of these 10 genes, 4 are targets of Polycomb Repressive Complex 1 (PRC1), 2 are targets of both PRC1 and PRC2 (“Both”) and 3 were not known or not direct targets of either.

| Gene Name | Gene Description | Gene Type | p.adj | log2FC | PRC Specificity |
| --- | --- | --- | --- | --- | --- |
| ADH1C | Alcohol Dehydrogenase 1C (Class I), Gamma Polypeptide | protein-coding | 0.046 | 1.16 | Not Known |
| DGCR11 | DiGeorge Syndrome Critical Region Gene 11 | lncRNA | 0.048 | 1.68 | PRC1 |
| IGFL4 | IGF Like Family Member 4 | protein-coding | 0.001 | 1.11 | Not Known |
| LINC00266-1 | Septin 14 Pseudogene 20 (SEPTIN14P20) | pseudogene | 0.025 | 1.54 | Not Known |
| MT1E | Metallothionein 1E | protein-coding | 1.3E-05 | 1.81 | Both |
| MT1G | Metallothionein IG | protein-coding | 1.44E-15 | 2.17 | PRC1 |
| MT1H | Metallothionein 1H | protein-coding | 0.0007 | 4.17 | PRC1 |
| SPP1 | Secreted Phosphoprotein 1 | protein-coding | 0.0006 | 2.01 | Not known |
| TNFRSF11B | TNF Receptor Superfamily Member 11b | protein-coding | 3.2E-18 | 1.16 | PRC1 |
| TNFSF12-TNFSF13 | Tumor Necrosis Factor (Ligand) Superfamily, Member 12-Member 13 | protein-coding | 3.72E-07 | 6.48 | Both |

**Table S3:**
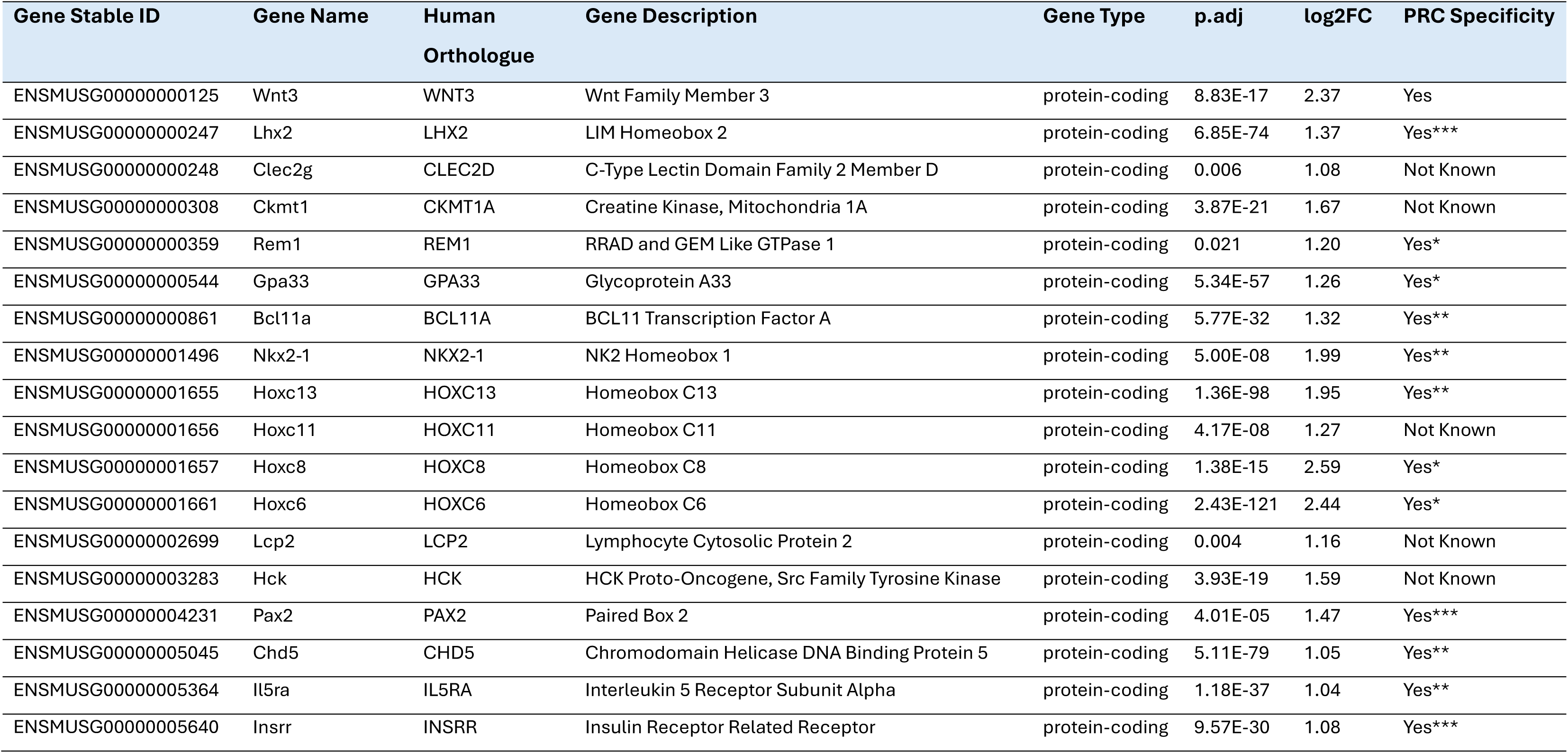
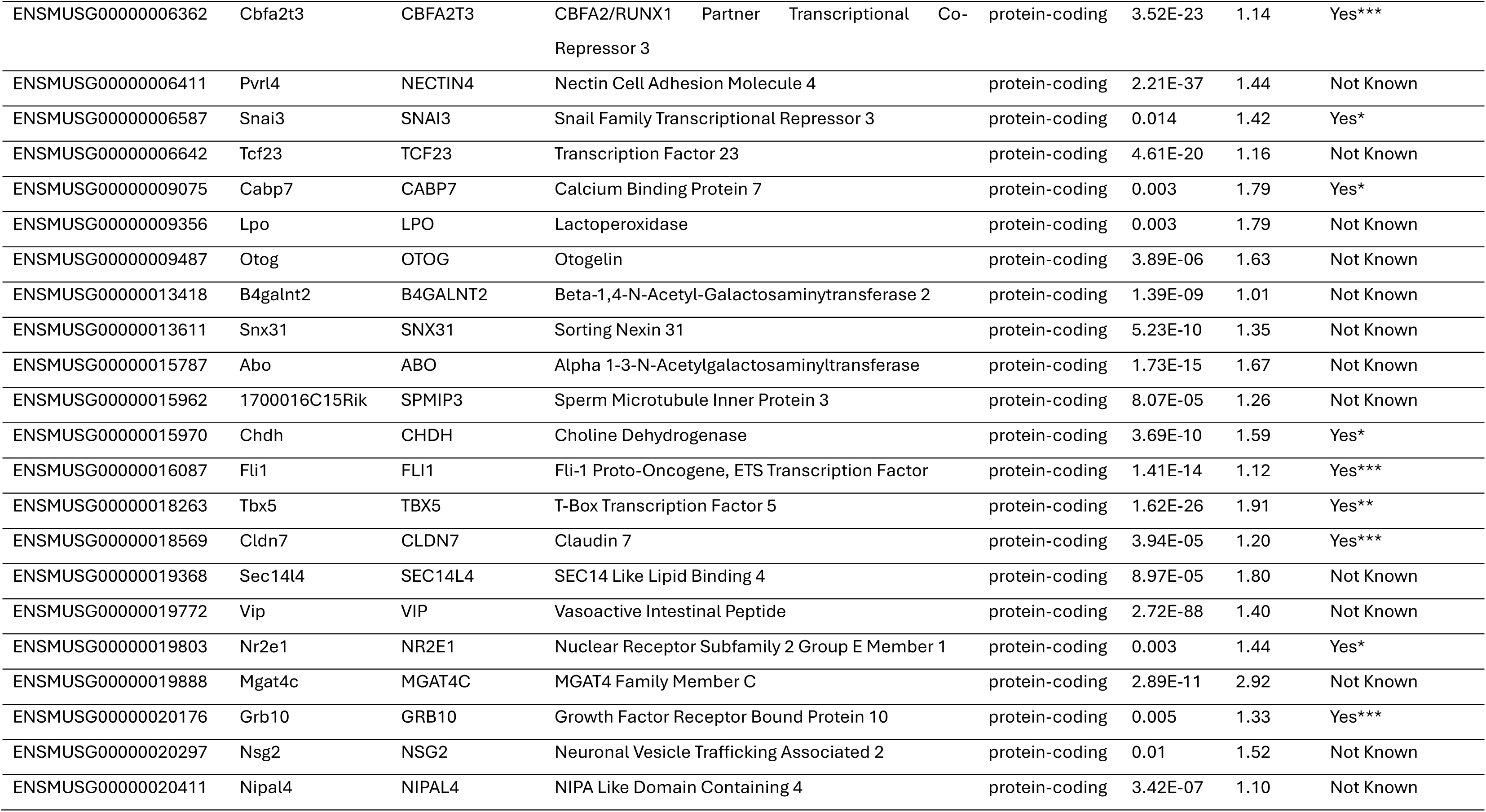
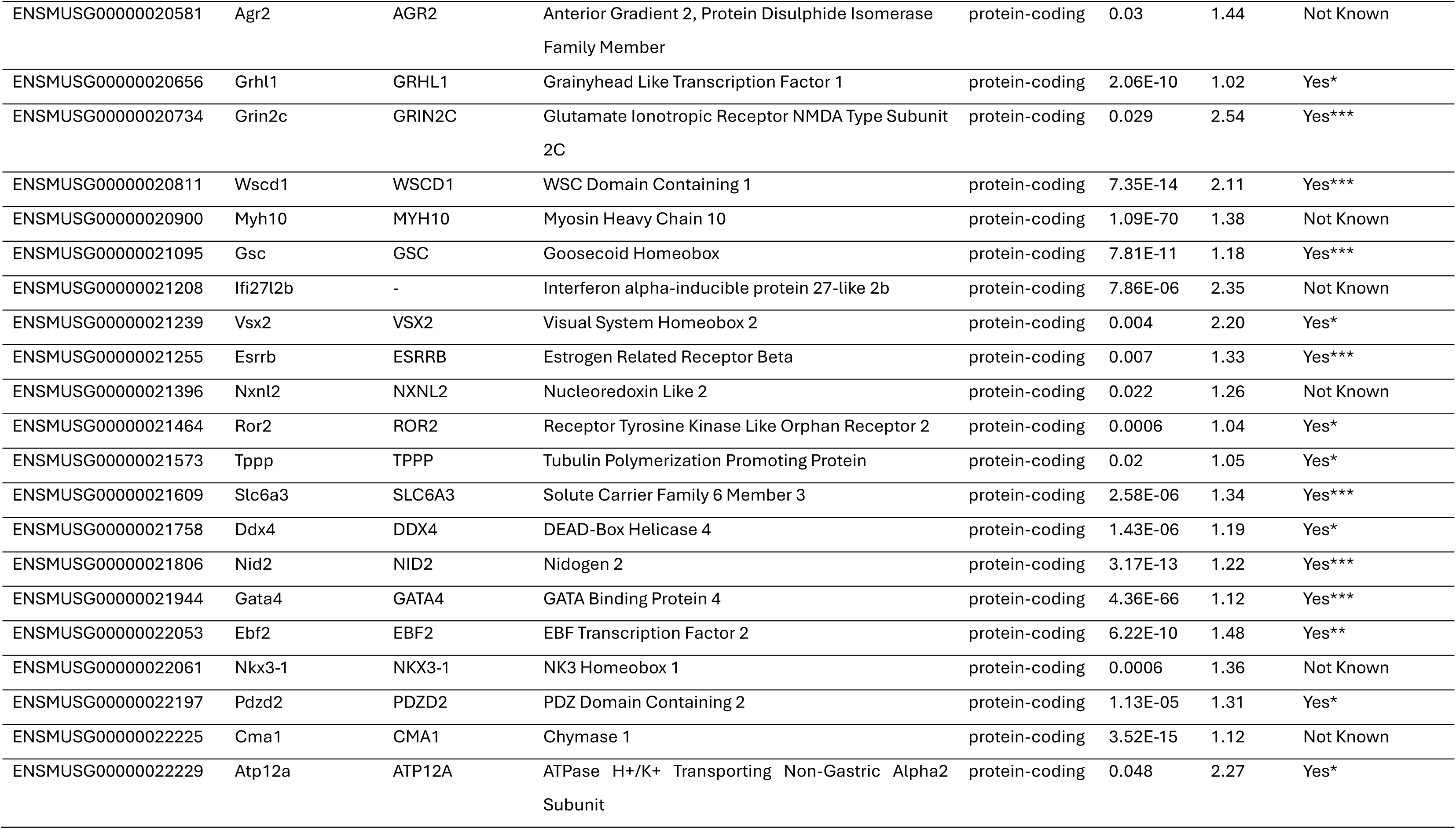
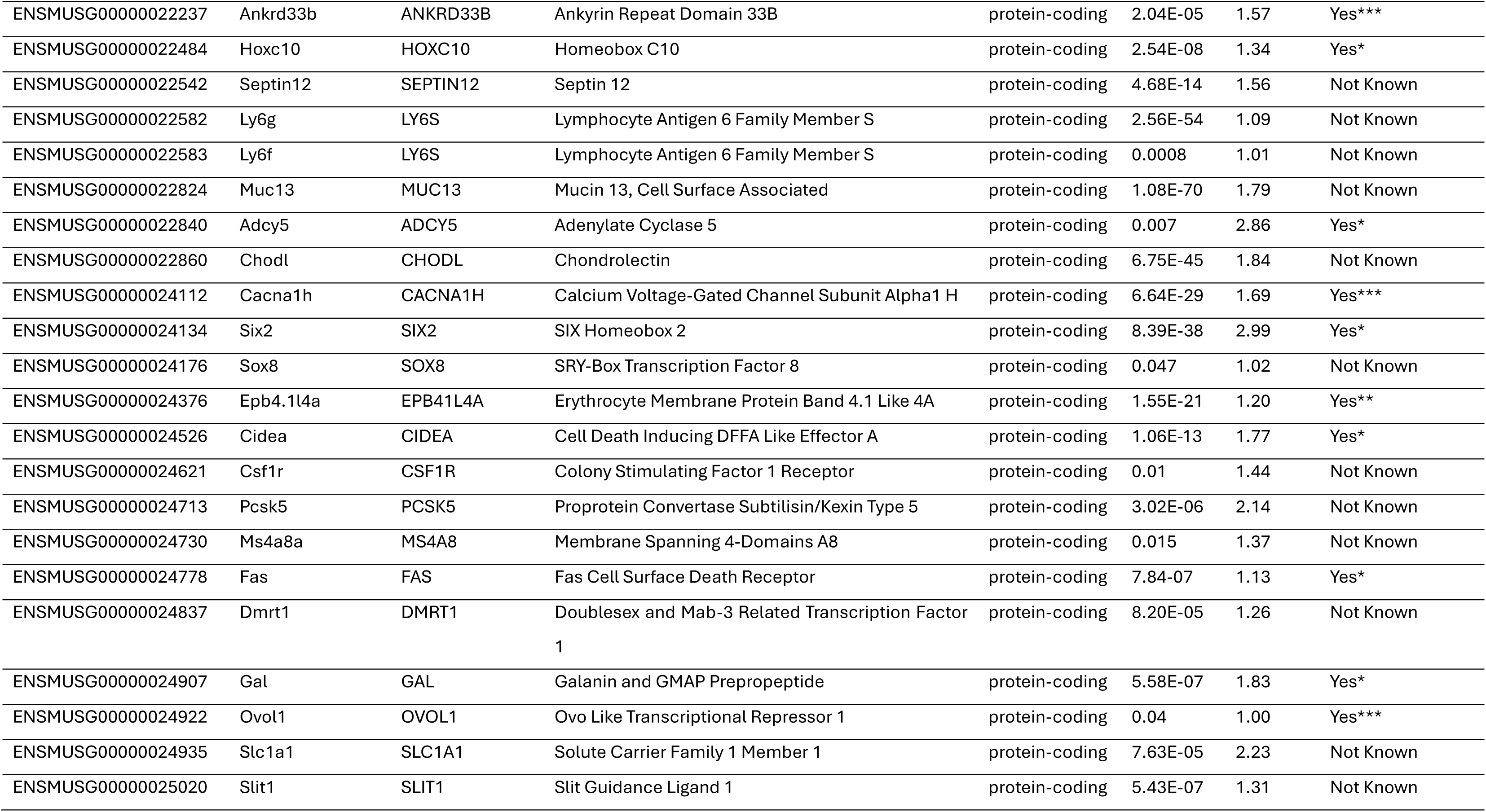
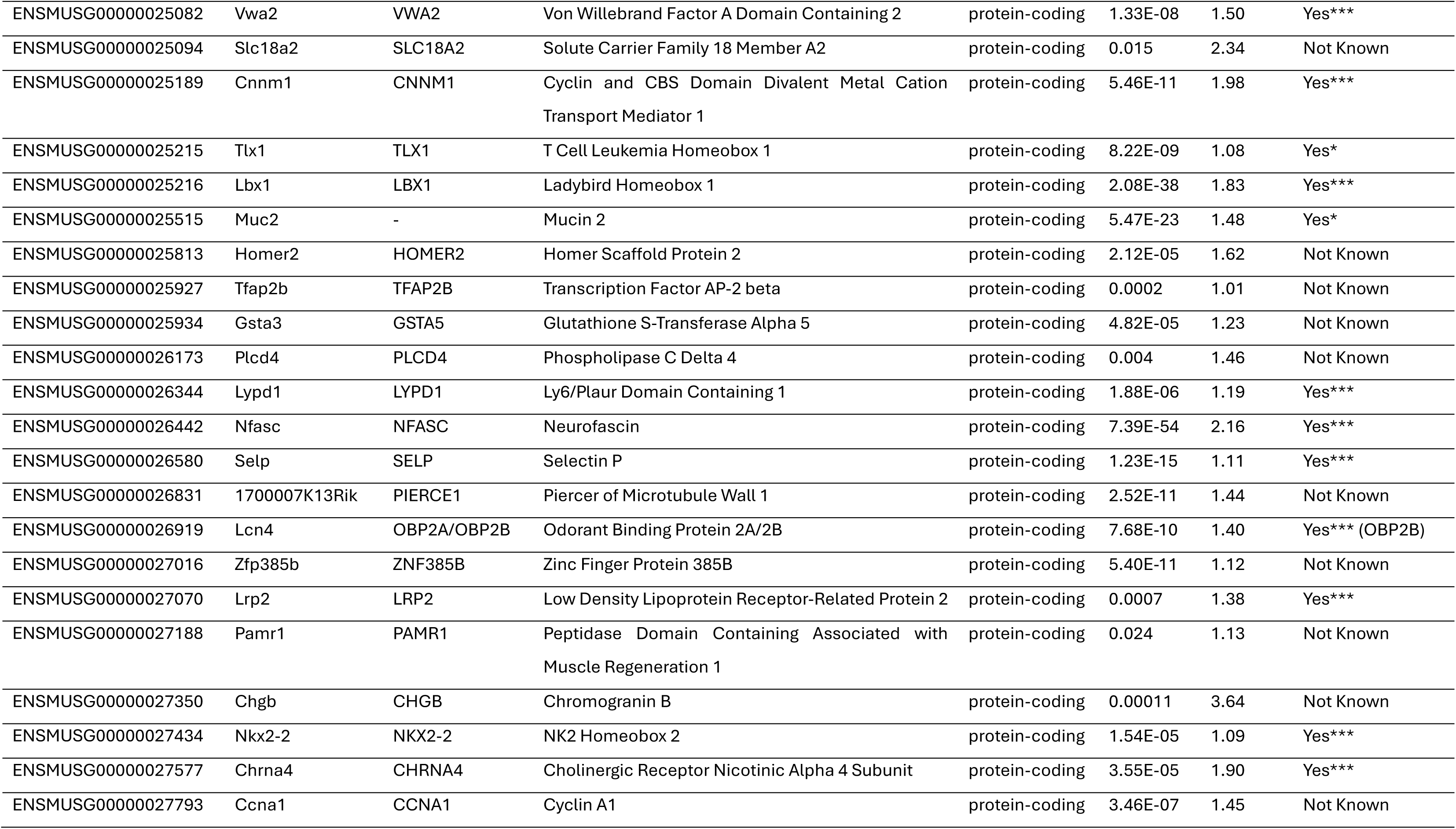
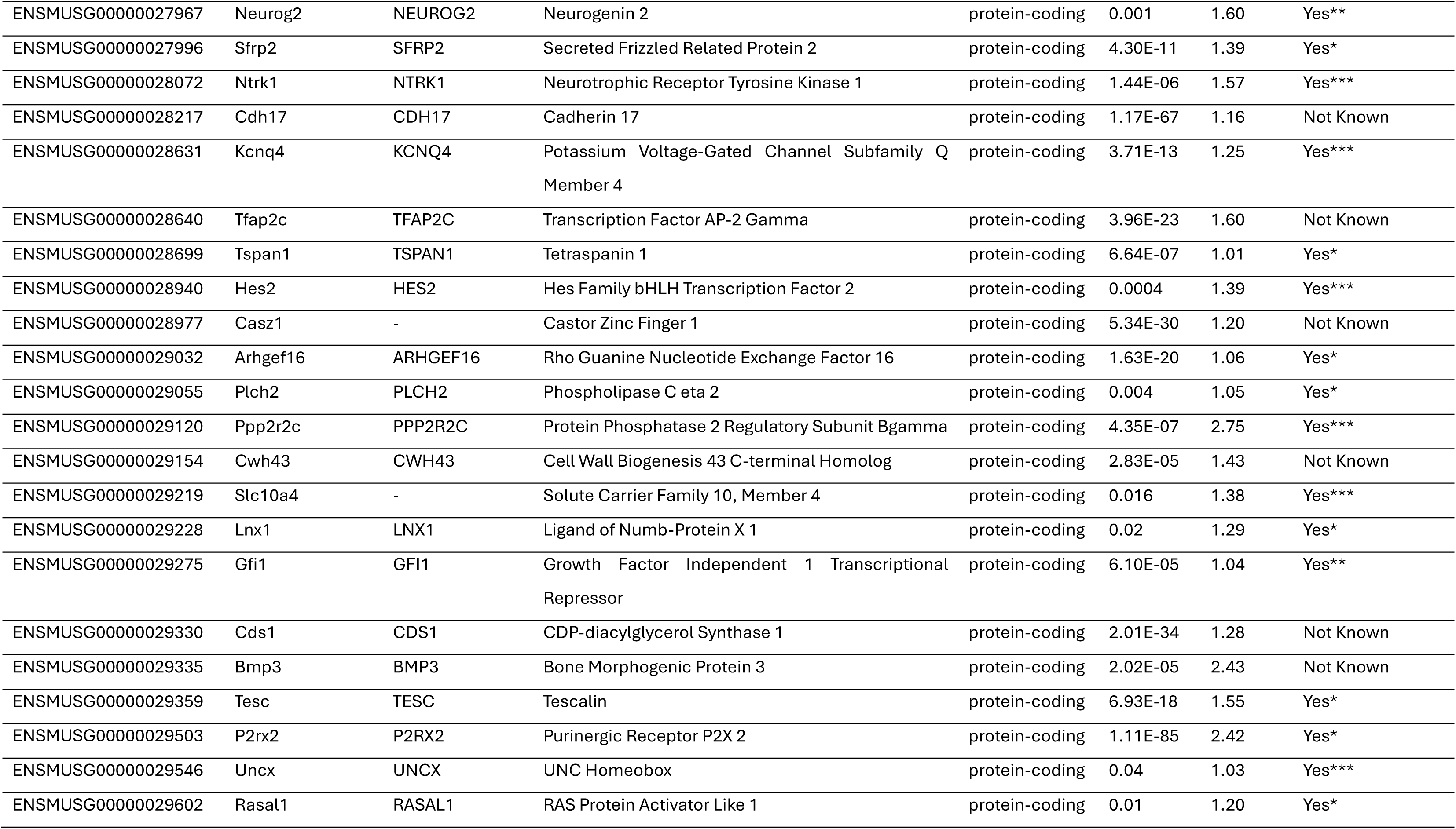
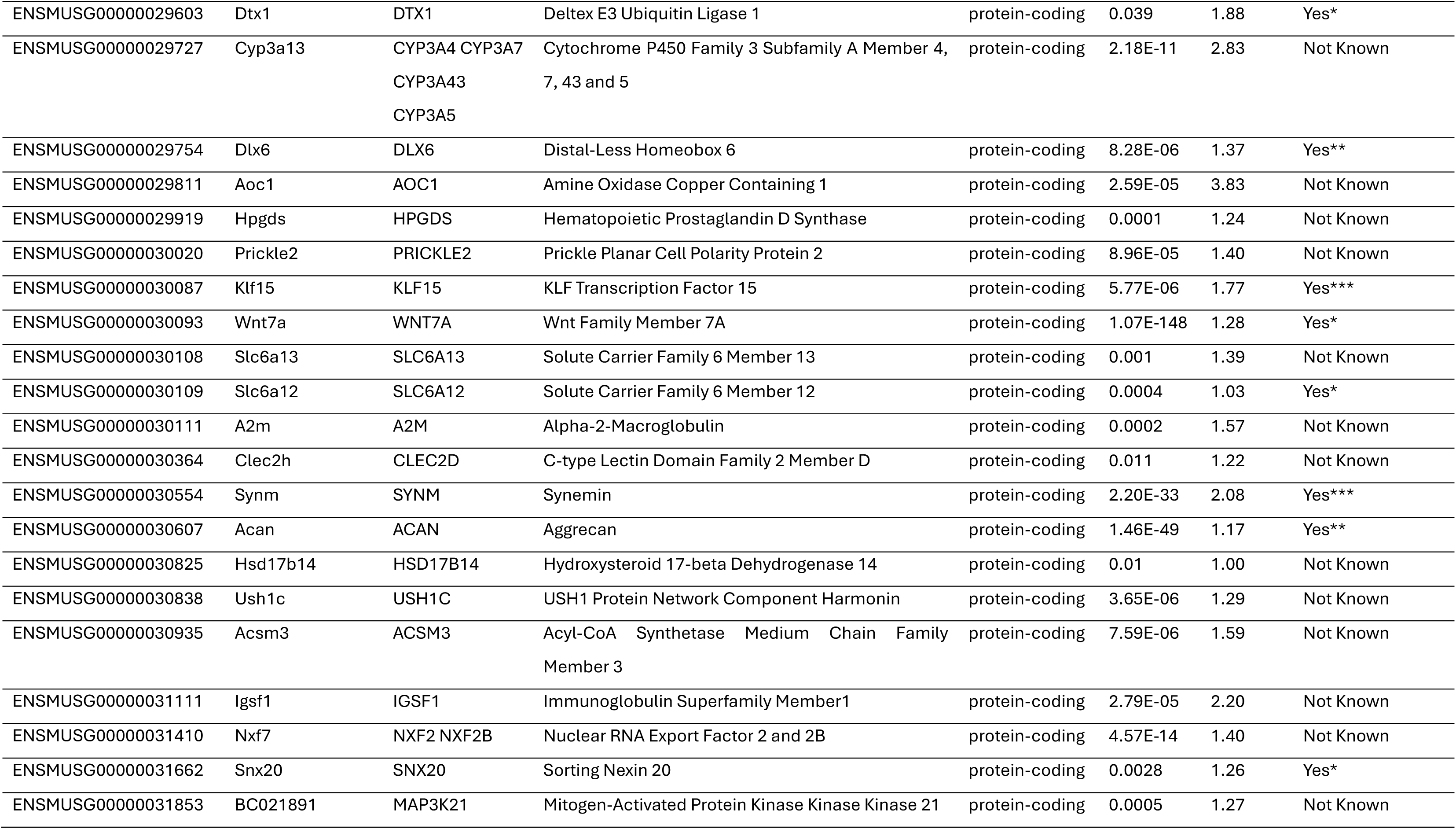
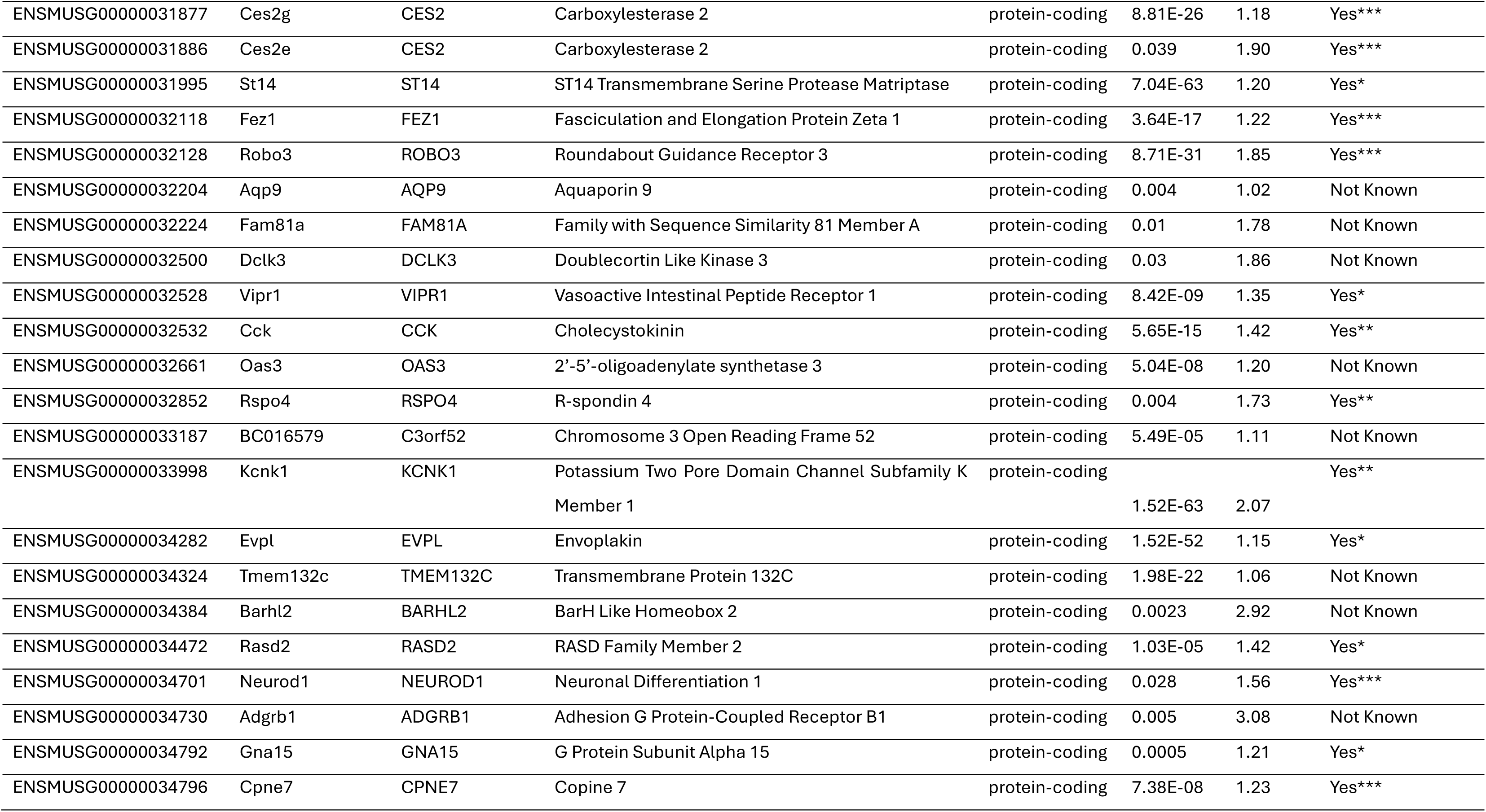
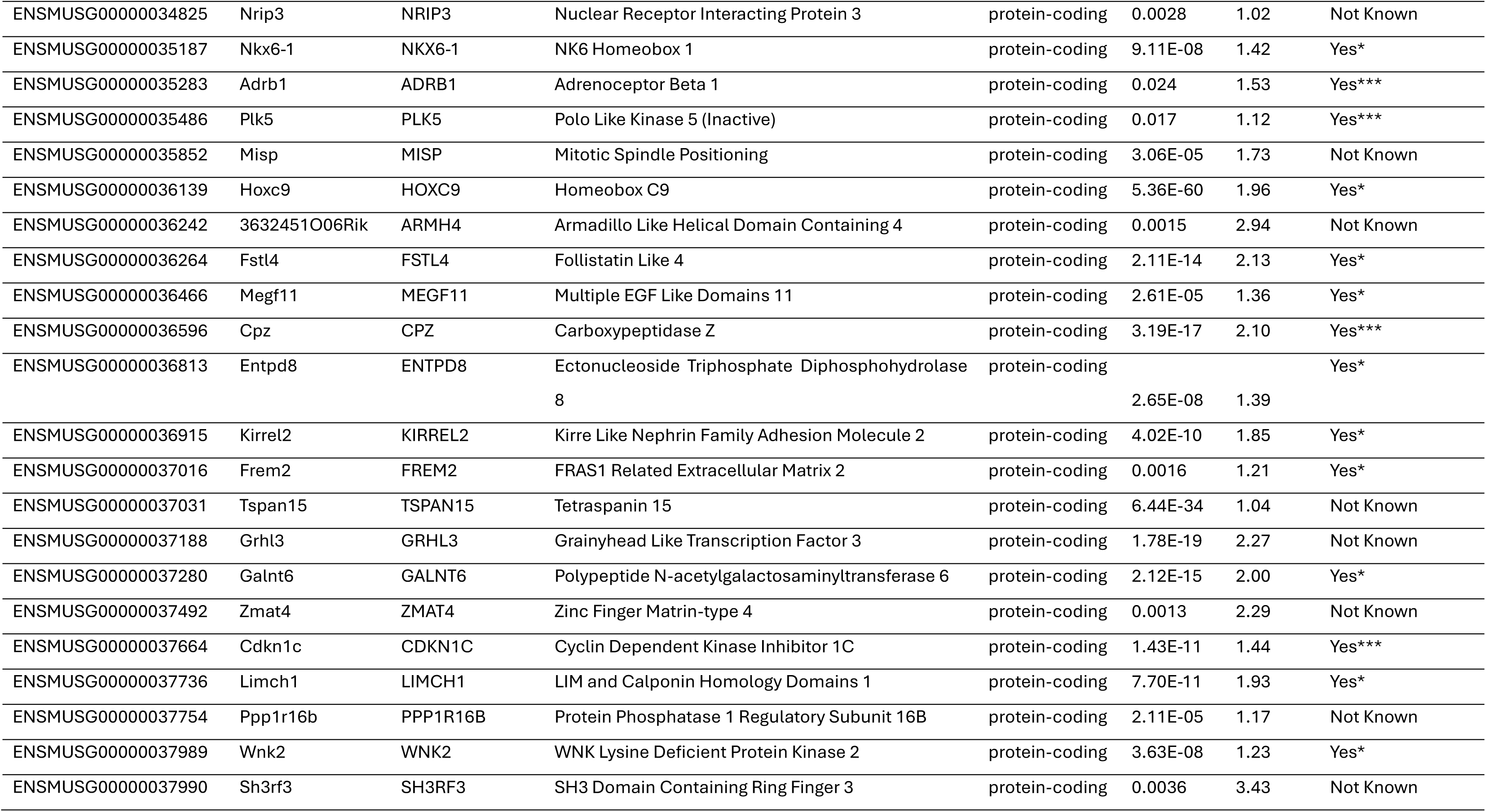
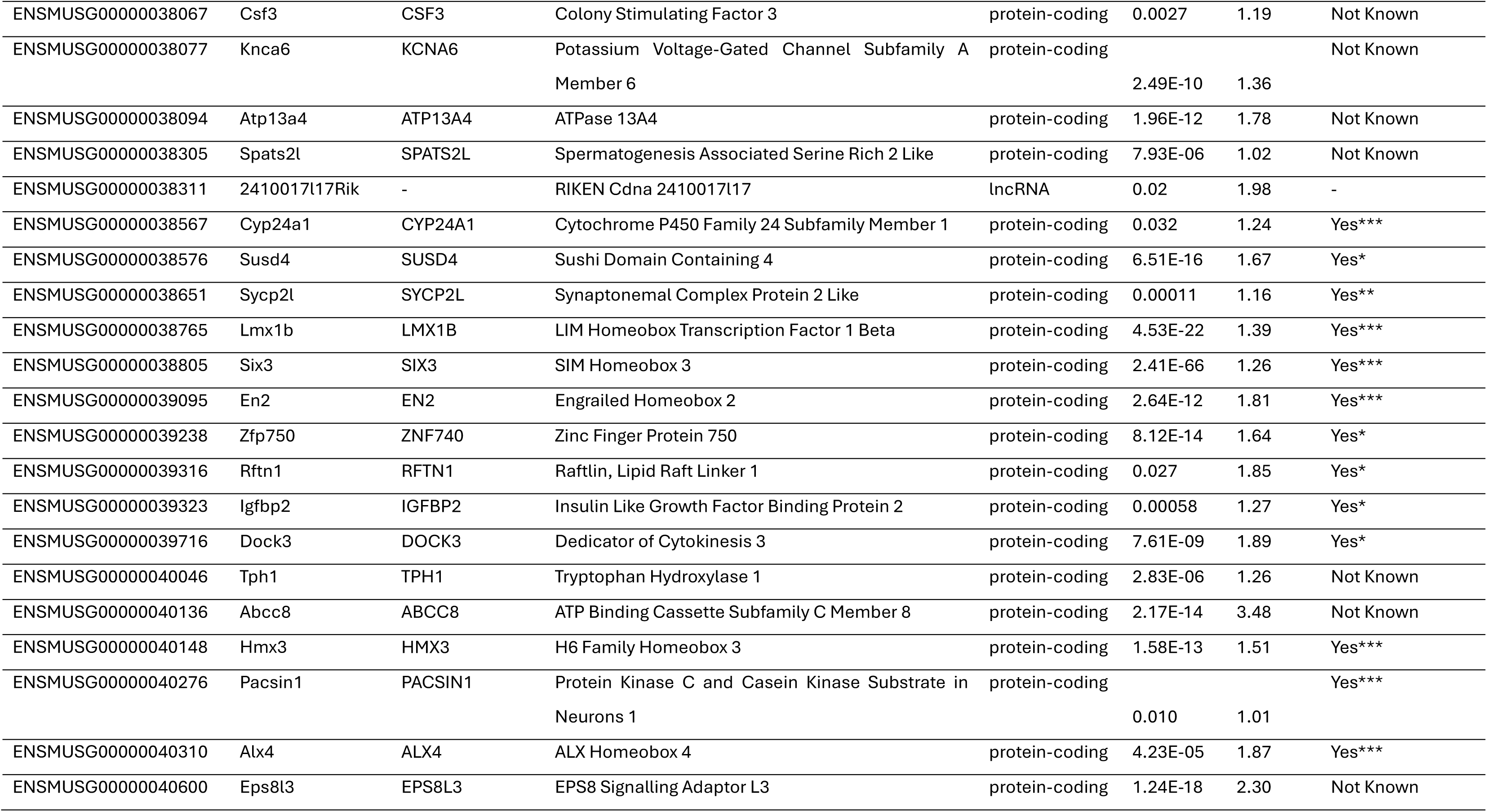
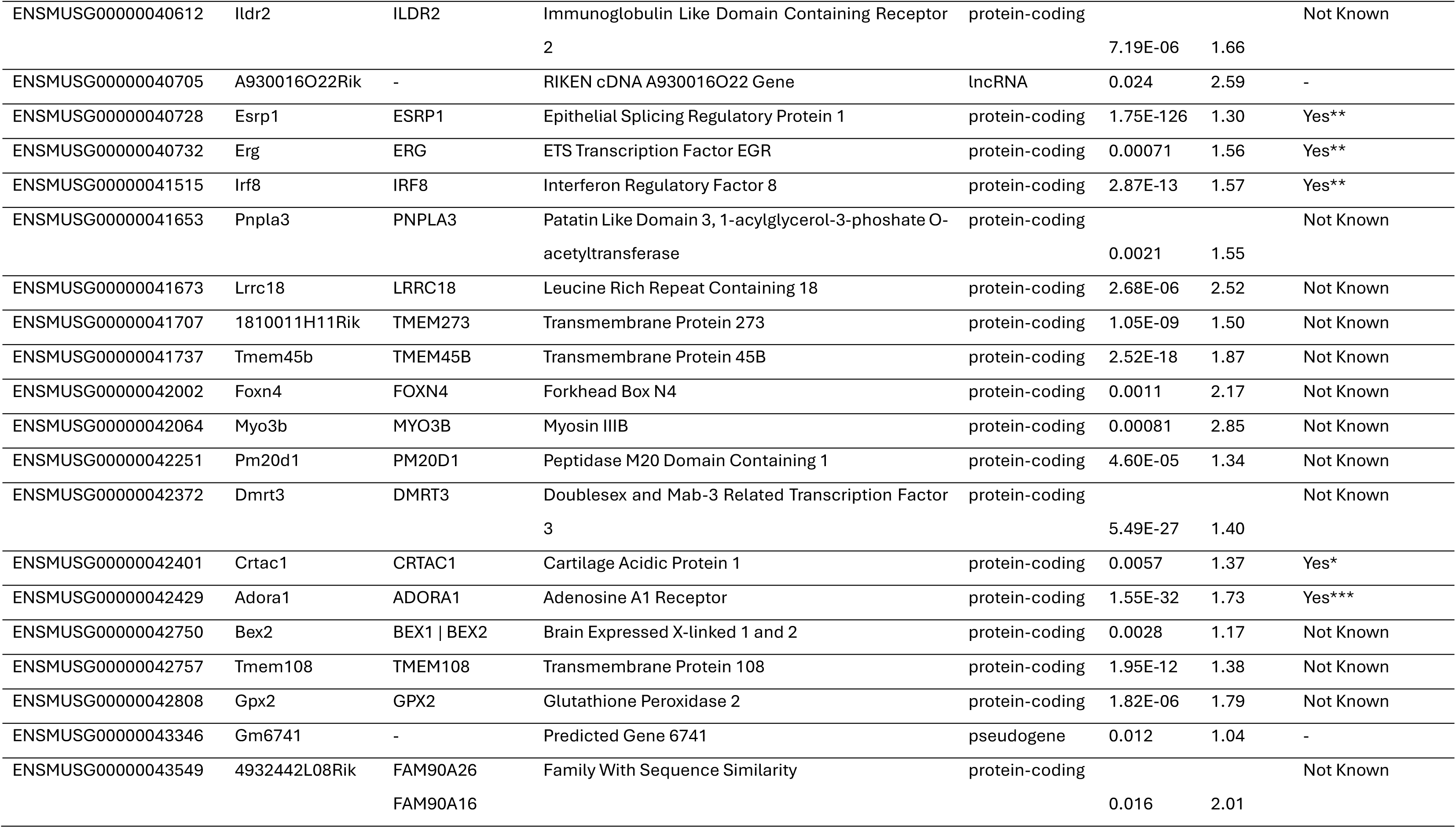
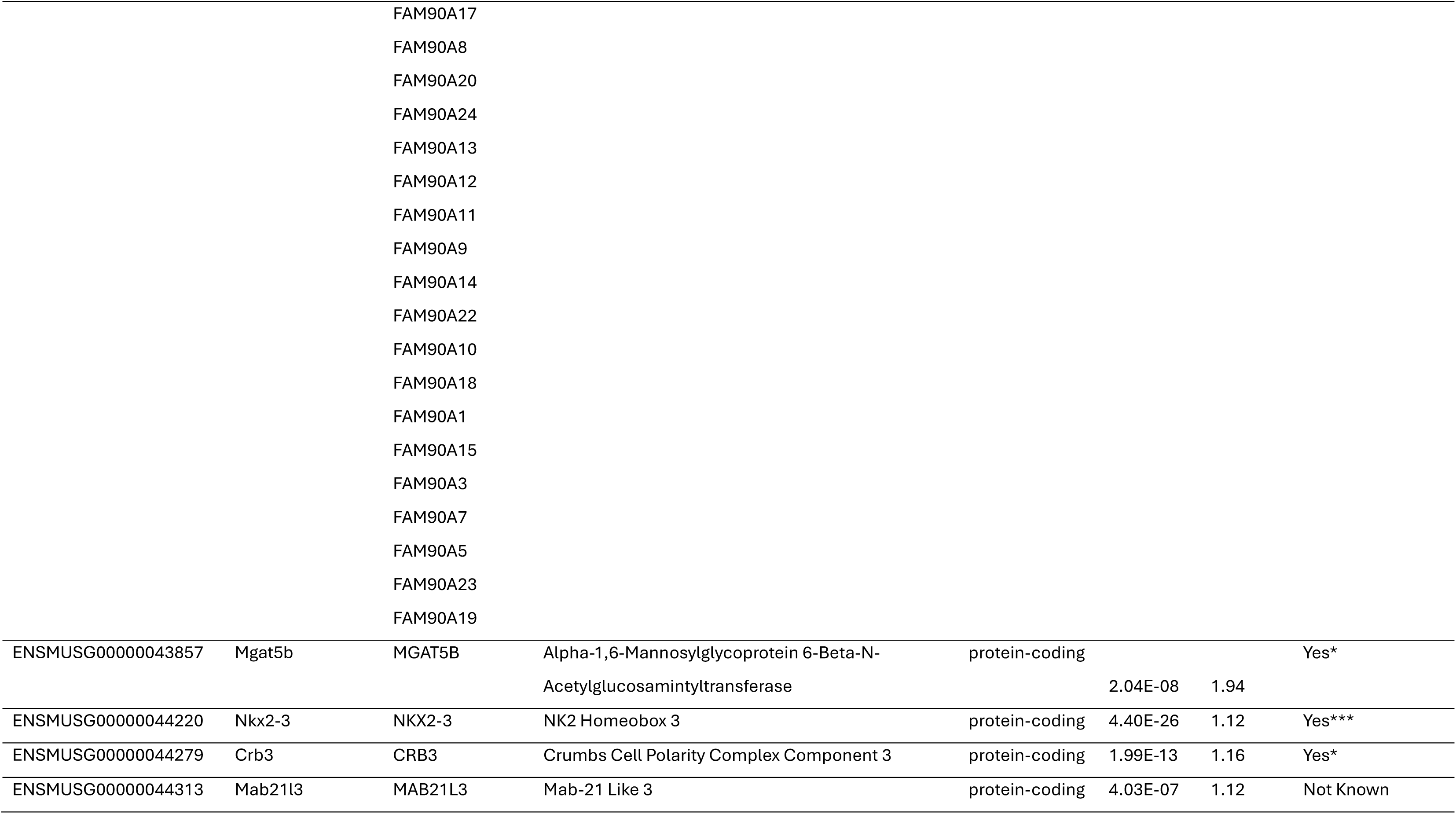
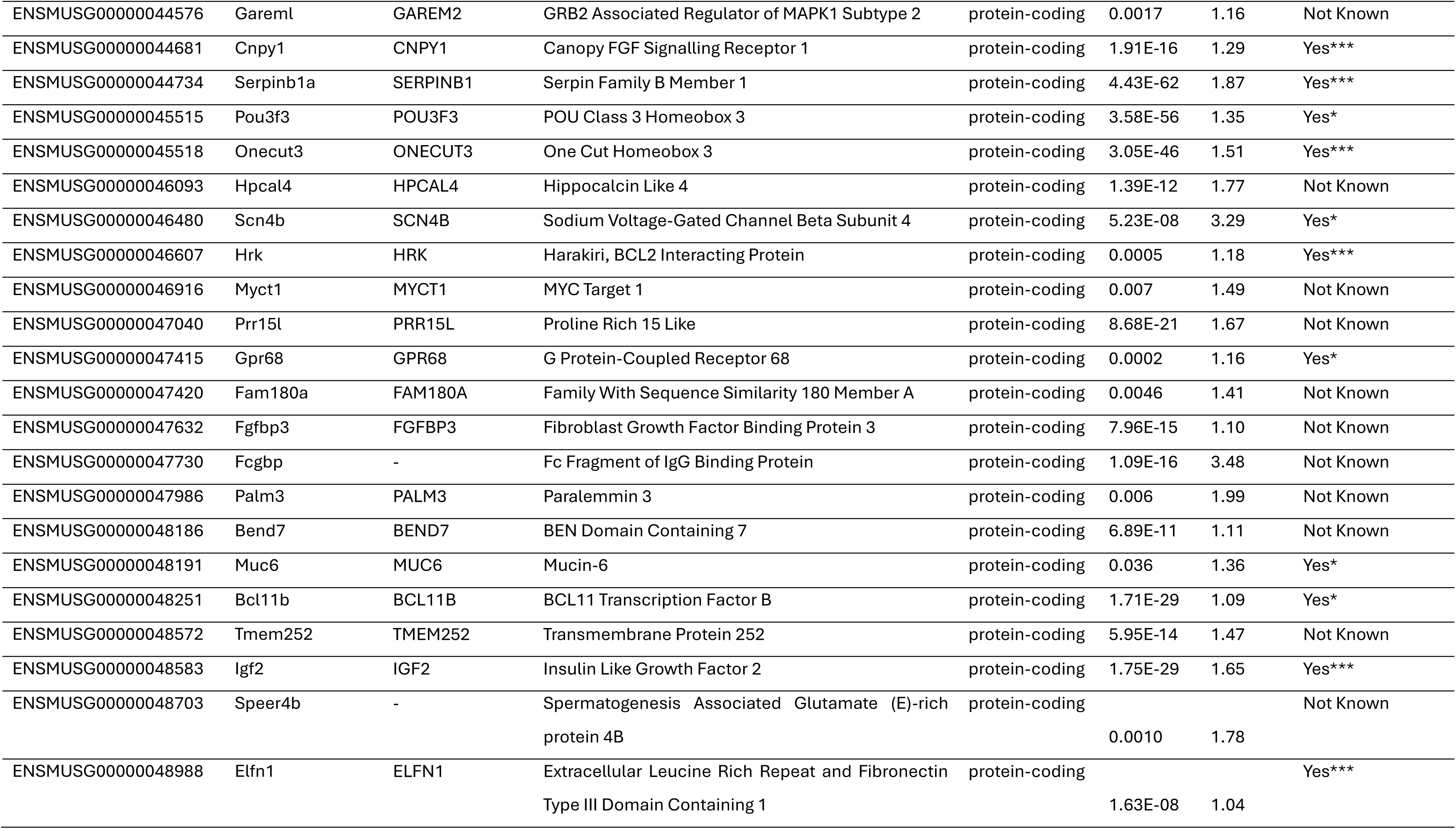
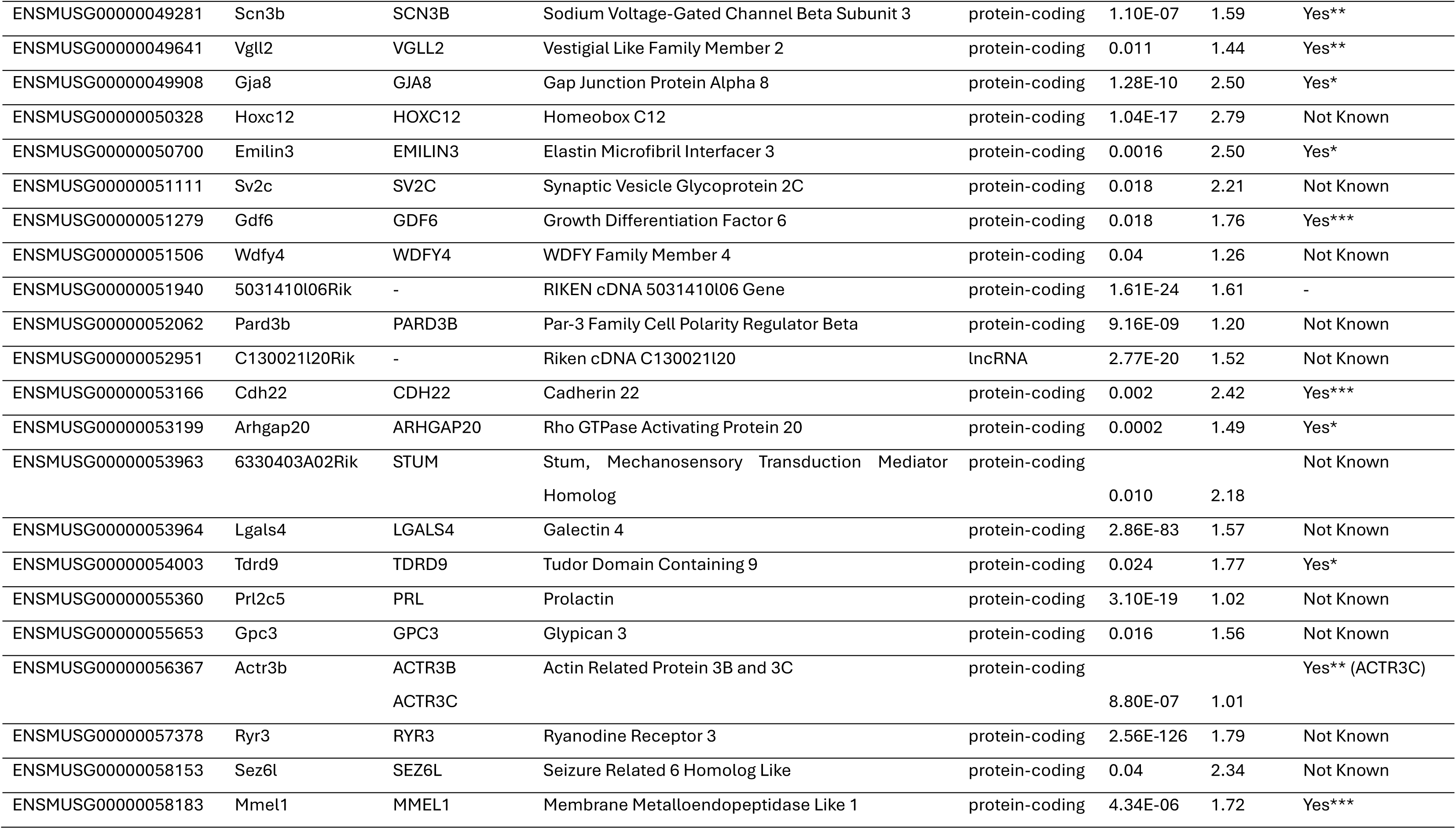
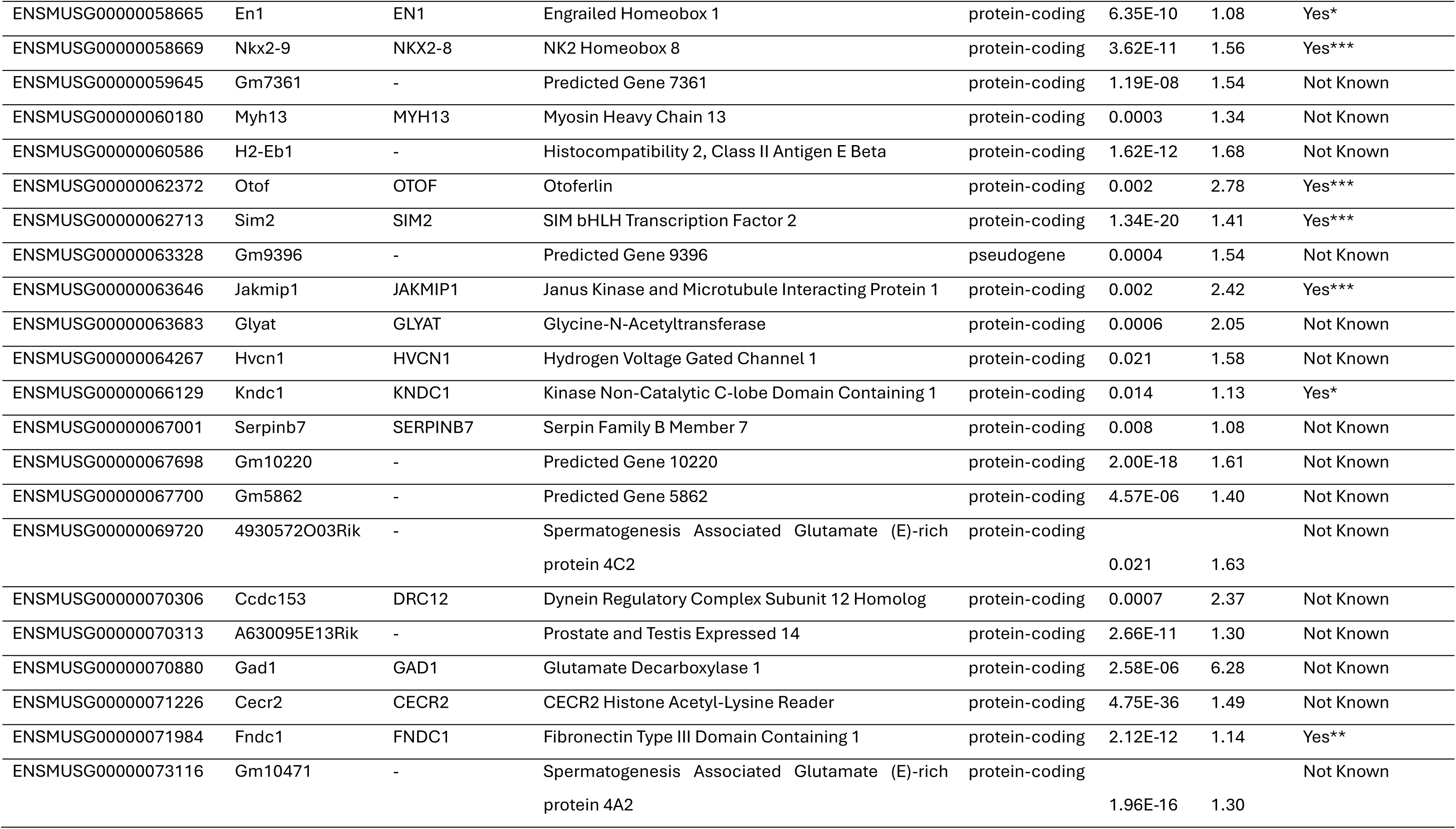
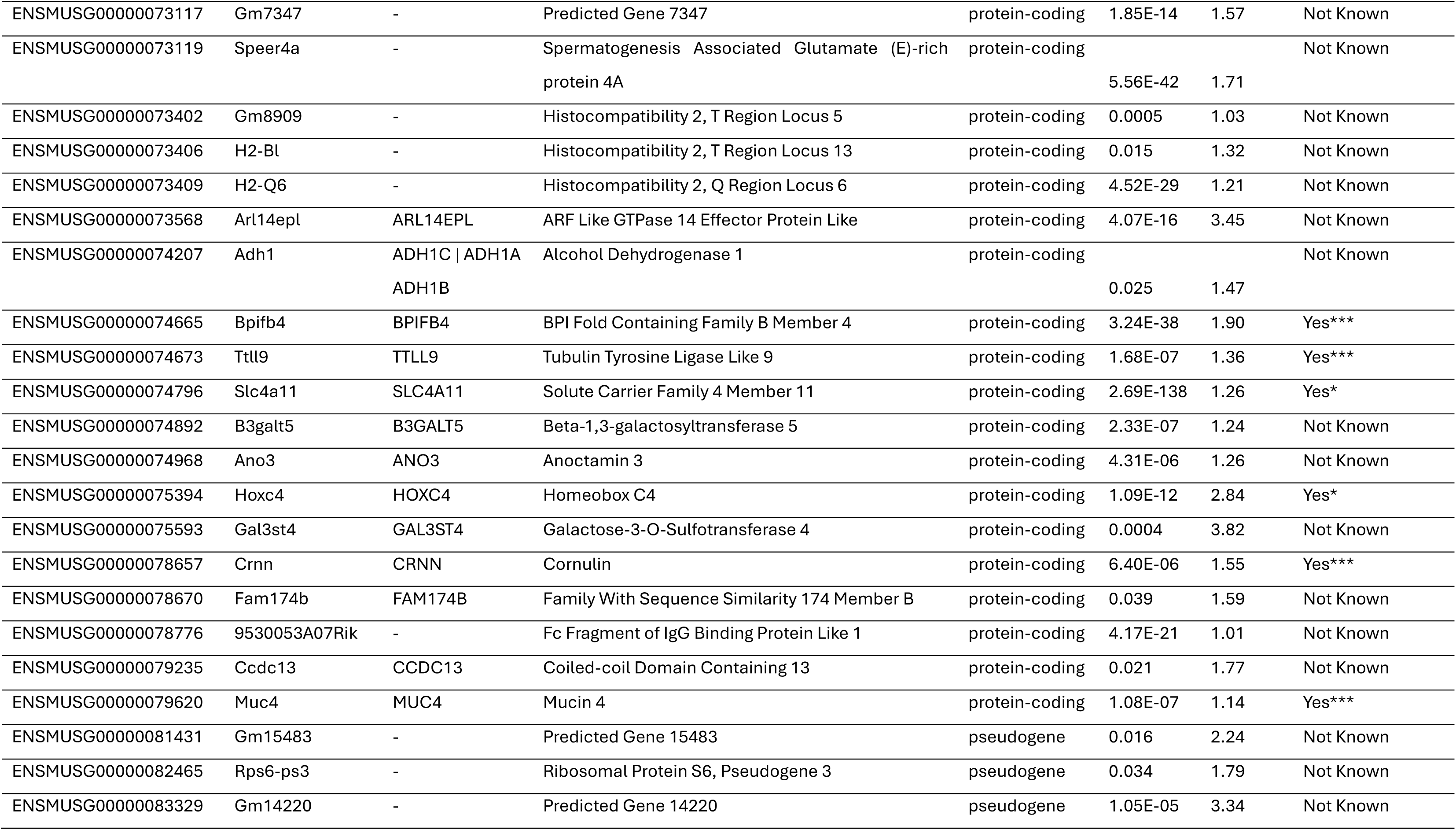
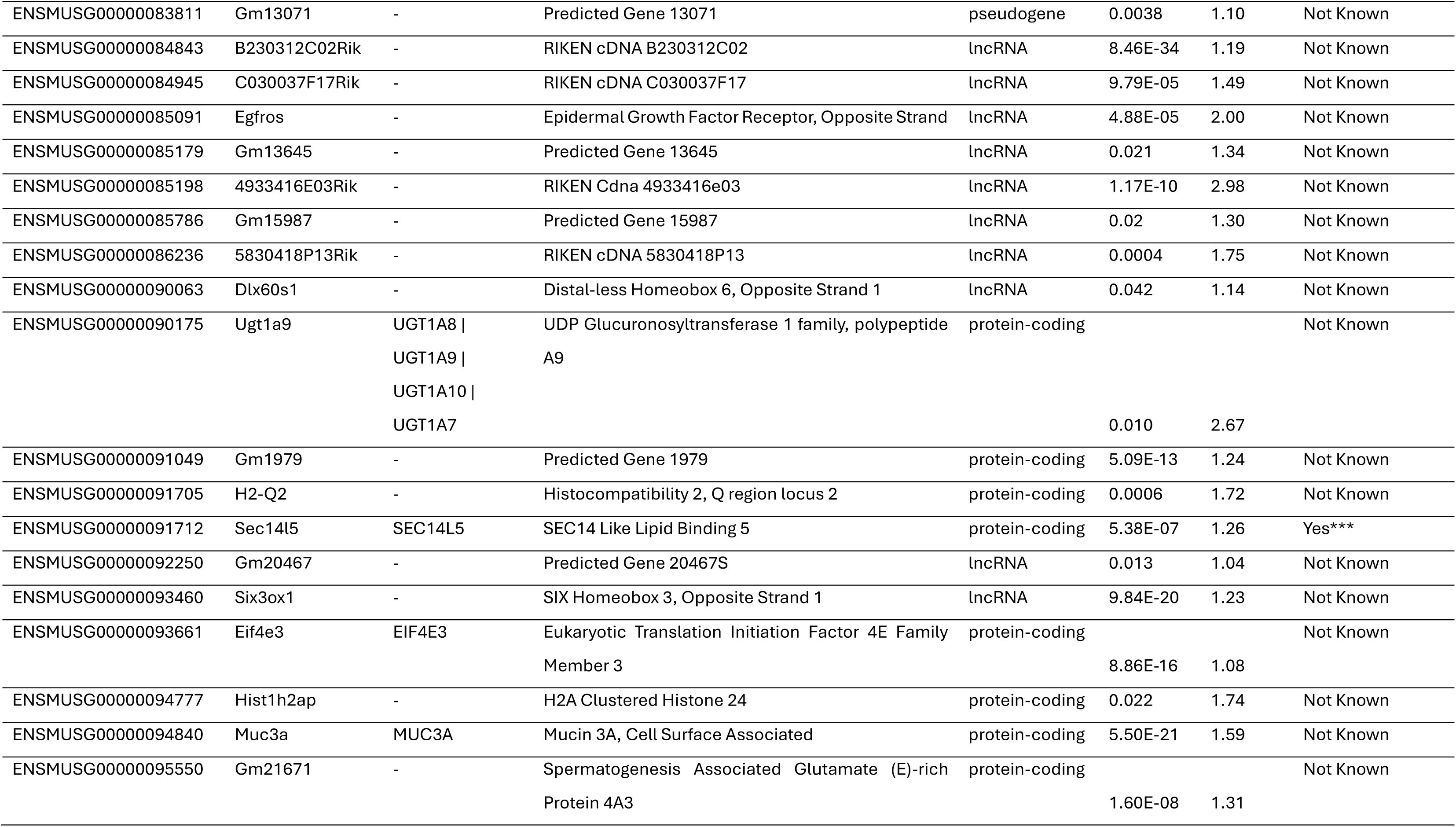
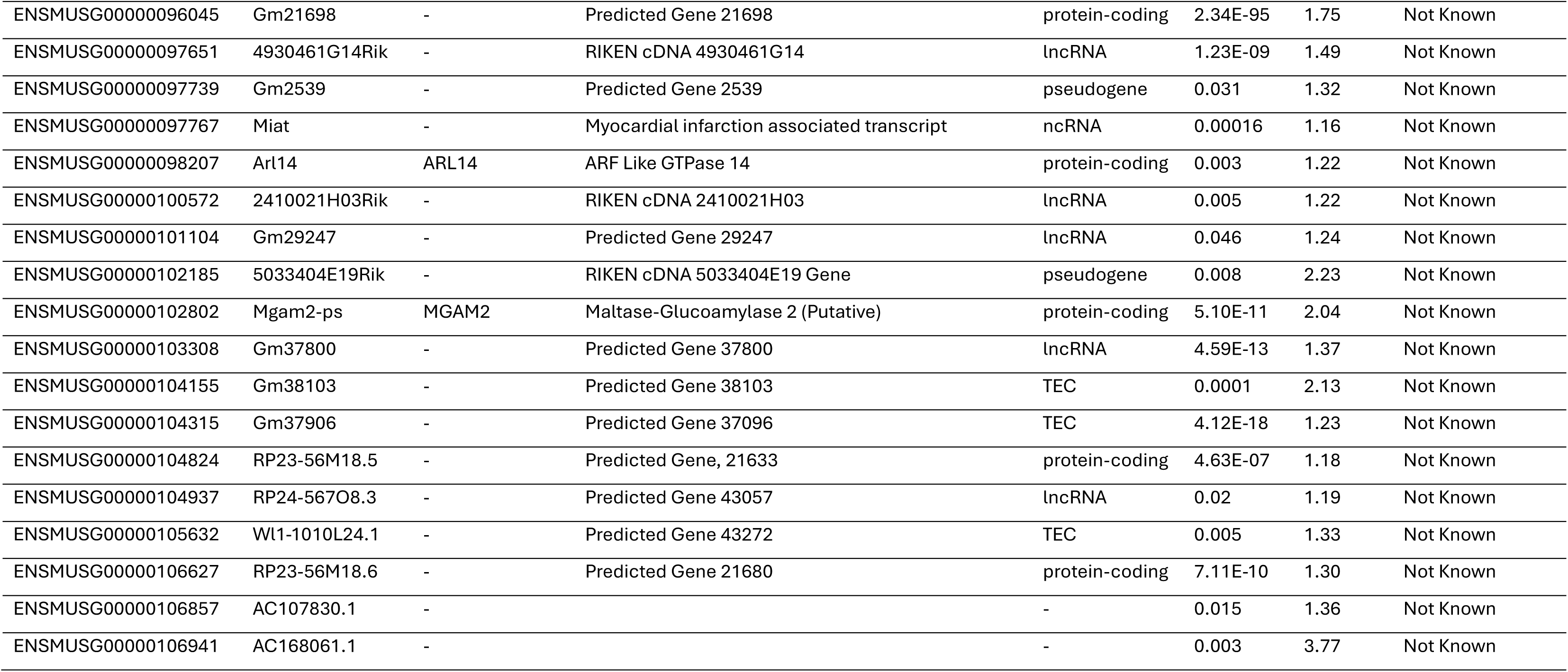
Preferentially ORIC-944 Dependent Genes (PODs) identified from DESeq2 analysis of ST4787. DESeq2 analysis of differential gene expression between tazemetostat and ORIC-944 in ST4787 identified 362 genes that were preferentially ORIC-944 dependent genes (PODs) as well as their p adjusted value (p.adj) and log2FoldChange (log2FC). This identified 330 protein coding, 18 lncRNA, 8 pseudogenes and 3 TEC genes. Additionally, using data obtained from DU145 scramble cells^[33]^, out of these 363 genes, 67 are targets of Polycomb Repressive Complex 1 (PRC1), 19 were targets of PRC2 (PRC2), and 70 were targets of both PRC1 and PRC2 (Both) and 191 were not known or not direct targets of either.

